# Human saphenous vein provides a unique source of anti-calcific pericytes for prosthetic cardiac valve engineering

**DOI:** 10.1101/2020.07.14.202846

**Authors:** Eva Jover, Marco Fagnano, William Cathery Meng, Sadie Slater, Emanuela Pisanu, Yue Gu, Elisa Avolio, Domenico Bruno, Daniel Baz-Lopez, Ashton Faulkner, Michele Carrabba, Gianni Angelini, Paolo Madeddu

## Abstract

**Aims:** Tissue engineering seeks to improve the longevity of prosthetic heart valves, but the cell source of choice has yet to be determined. This study aimed to establish a mechanistic rationale supporting the suitability of human adventitial pericytes (APCs).

**Methods and Results:** **Antigenically** APCs were immunomagnetically sorted from saphenous vein leftovers of patients undergoing coronary artery surgery and antigenically characterized for purity. Unlike bone marrow-derived mesenchymal stromal cells (BM-MSCs), APCs were resistant to osteochondrogenic induction by high phosphate (HP), as assessed by cytochemistry and expression of osteogenic markers. MiR-132 is natively expressed by APCs, with copy numbers being enhanced by HP stimulation. *In silico* bioinformatic analysis, followed by luciferase assays in HEK293 cells and miR-132 titration using agomiR and antagomiR in APCs, demonstrated that several osteochondrogenic genes were negatively regulated by miR-132. Among these, the glycolytic marker GLUT1 was downregulated in HP-stimulated APCs. In contrast to BM-APCs, APCs showed no increase in glycolysis under HP. Interestingly, incubation with APC-derived conditioned medium conferred swine cardiac valves with resistance to osteogenic transformation by HP; whereas, conditioned media from miR-132-knocked-down APCs failed to prevent the expression of these markers. Finally, we demonstrated the feasibility of using APCs to engineer bovine pericardium patches. APCs proliferate in the patch and secrete factors able to attract aortic endothelial cells under HP.

**Conclusions:** Human APCs are resistant to calcification compared with BM-MSCs and convey the anti-calcific phenotype to heart valves through miR-132. These findings may open new important avenues for prosthetic valve cellularization.

## Introduction

Calcific valvular heart disease (VHD) represents the third most common cardiovascular pathology in adults after hypertension and coronary artery disease.^1, 2^ Aortic valve disease alone affects ~2% of the population over 65 years, and is a major cause of morbidity and mortality in the elderly.^3, 4^ Surgical valve replacement remains one of the main therapeutic solutions to treat VHD and accounts for more than 20% of all cardiac surgeries.^5^

Over the past decades, the clinical outcome of patients undergoing valve substitution has been significantly improved thanks to more frequent use of biological prostheses and better control of risk factors and complications. Biological prostheses, however, undergo calcific degeneration, ultimately requiring reinterventions after 10-15 years from implantation.^6, 7^

Tissue engineering of heart valves (TEHV) promises to overcome the current limitations by constructing a living valvular substitute capable of physiological remodelling through exogenously implanted cells (reviewed in ^8^). Among the different cells proposed so far, bone marrow-mesenchymal stromal cells (BM-MSCs) remain the ‘gold standard’;^9^ though modest results have been reported in recent clinical trials.^10–12^ Adventitial pericytes (APCs) represent a newly characterized clonogeneic stromal cell population,^13, 14^ which reportedly surpassed MSCs in terms of purity and therapeutic potential,^15, 16^ and was also suitable for xenograft cellularization.^17^ After selective immunomagnetic sorting from vascular tissue, APCs can be expanded through different passages whilst maintaining typical pericyte (PDGFRβ and CSPG4/NG2) and mesenchymal markers (vimentin, desmin, CD90, CD44, CD29, CD105, CD49a, CD49b, CD13, CD59, and CD73).

In models of ischemia, APC transplantation exerted remarkable therapeutic benefit by promoting reparative angiogenesis and inhibiting fibrosis.^15, 16^ MicroRNA-132 (miR-132) emerged from preclinical studies as one of the APC secreted factors responsible for main therapeutic actions.^16^ The role of miR-132 in cardiovascular disease remains controversial. Transgenic overexpression of the miR-212/132 cluster results in pathological cardiac remodelling;^18^ while other reports suggest that miR-132 promotes vascular growth.^19^ These differences could be attributed to additional auxiliary binding of miR-212 to its targets and to the magnitude and context of miR-132 expression. Seminal evidence indicates that miR-132 directly targets the expression of MeCP2 (validated target) and calumenin (predicted target), which modulate osteogenesis by interfering respectively with the Wnt signalling pathway and vitamin K-dependent γ-carboxylation of matrix Gla protein.^20–22^ Nonetheless, it remains unknown whether APCs could be resistant to calcification *via* a mechanism involving miR-132 signalling.

The aim of present study was three-fold: (1) to compare the susceptibility of human APCs and BM-MSCs to undergo osteochondrogenic transition following exposure to HP, (2) to determine the role of miR-132 in the above phenomena, and (3) to assess the suitability of APCs for cellularization of FDA-approved bioprosthetic material.

## Methods

An extended version of material and methods is provided as online supplementary material

### Cell isolation and culture

Studies were performed on leftover material, according to the ethical principles recorded in the 1964 Declaration of Helsinki, and covered by the following Research Ethics Committee approvals (06/Q2001/197 covering use of APCs, and 14/SW/1083 and 14/WA/1005 covering use of BM). All recruited subjects provided informed written consent. Human APCs were isolated and expanded from saphenous vein leftovers of patients undergoing coronary artery bypass graft surgery (performed at the Bristol Royal Infirmary Hospital, UK), as previously published.^15, 16^ BM was obtained from femur heads of patients undergoing hip replacement surgery at the Avon Orthopaedic Centre (Southmead Hospital, UK) by using the Ficoll stratification method (see **Supplementary Material**). **Table 1** and **2** summarizes clinical and demographic data of APC and BM-MSC donors.

**Table 1.** Clinical and demographic data of APCs donors

| **Baseline characteristics** |  | **n = 20** |
| --- | --- | --- |
| Male sex, n (%) |  | 17 (80.95%) |
| Age, median [IQR] (n) |  | 71.50 [60.25-75.00] (18) |
| Hypertension, n (%) |  | 16 (80%) |
| Diabetes mellitus, n (%) |  | 2 (10%) |
| Hyperlipidaemia, n (%) |  | 16 (80%) |
| Body mass index (kg/m2), median [IQR] (n) |  | 28.90 [25.50-33.23] (18) |
| Smoking habit, n (%) |  |  |
|  | Non-smoker | 7 (35%) |
|  | Previous | 9 (45%) |
|  | Current | 2 (10%) |
|  | Unknown | 2 (10%) |
| Previous Myocardial Infarction |  |  |
| Coronary Artery Disease, n (%) |  |  |
|  | 0 vessels affected | n/a |
|  | 1 vessel affected | n/a |
|  | 2 vessels affected | 2 (10%) |
|  | 3 vessels affected | 16 (80%) |
|  | Unknown | 2 (10%) |
| Angina classification (CSS score), n (%) |  |  |
|  | Asymptomatic | 1 (5%) |
|  | I | 4 (20%) |
|  | II | 3 (15%) |
|  | III | 7 (35%) |
|  | IV | 3 (15%) |
|  | Unknown | 2 (10%) |
| NYHA classification, n (%) |  |  |
|  | I | 6 (30%) |
|  | II | 6 (30%) |
|  | III | 6 (30%) |
|  | Unknown | 2 (10%) |
| Left ventricular function, n (%) |  |  |
|  | Good (>50) | 12 (60%) |
|  | Moderate (30-50) | 6 (30%) |
|  | Low (<30) | n/a |
|  | Unknown | 2 (10%) |

**Table 2.** Clinical and demographic data of BM-MSC donors

| Baseline characteristics | (n =4) |
| --- | --- |
| Male sex, n (%) | 2 (50%) |
| Age, median [IQR] (n) | 56.00 [41.50-75.75] |
| Hypertension, n (%) | 4 (100%) |
| Diabetes mellitus, n (%) | 0 (0%) |
| Hyperlipidaemia (total cholesterol > 5.2 mmol/L), n (%) | 1 (25%) |
| Body mass index (kg/m2), median [IQR] (n) | 31.70 [28.9-41.40] (4) |
| Smoking habit, n (%) | 0 (0%) |
| Other pathologies | None described |

HEK293 cells (CRL-11268™, ATCC^®^. Gaithersburg, Maryland, US) were cultured in DMEM high glucose (31966-021, Gibco™, Thermo Fisher, UK), supplemented with 10% FBS (16140-071, Gibco™, Thermo Fisher, UK), and used in luciferase assays to validate miR-132 binding to the *in silico* predicted sites on selected targets. Human aortic endothelial cells (AorECs) (C-12271, Promocell, UK) were cultured in complete EGM2 following manufacturer’s instructions.

### Osteogenesis

HP (2.6mM inorganic phosphate) was used as osteogenic stimuli on APCs and BM-MSCs at passage 3-6. Growing media were supplemented with 5 times less FBS and 2.6 mM HP buffer (Na_2_HPO_4_/NaH_2_PO_4_, pH 7.4). Previously published protocols were adapted,^22, 23^ and media were replaced every 3 days. Calcification endpoint was assessed by cell monolayer cytochemistry (Alizarin Red and von Kossa stainings), *o*-cresoftalein method (ab102505, abcam, UK) on 0.6N HCl hidroacidic extracts as previously published.^23, 24^ APCs and BM-MSCs from 4 donors each were assayed in technical triplicates and averaged for statistical analysis. RNA isolation and qPCR were performed to assess osteogenesis. Western blotting or enzyme-linked immunosorbent assays (ELISA) were used to validate gene expression studies.

### RNA isolation, RT and qPCR

Total RNA was isolated according to a standardized phenol-chloroform protocol, using Qiazol reagent and miRNeasy mini Kit (217004, QIAGEN, Germany), and reverse-transcribed into single-stranded cDNA, using a High Capacity RNA-to-cDNA Kit (4387406, Applied Biosystems™, Thermo Fisher, UK), or specific Taqman microRNA assay primers with a TaqMan^®^ MicroRNA Reverse Transcription Kit for the assessment of microRNAs (4366596, Applied Biosystems™, Thermo Fisher). Downstream qPCR amplifications of first-strand cDNA were performed using TaqMan^®^ Universal PCR Master Mix, no AmpErase^®^ UNG (4324018) or Power SYBR^®^ Green PCR Master Mix (4367659) (both from Applied Biosystems™, Thermo Fisher) in an Applied Biosystems QuantStudio 5 Real-Time PCR System. The relative expression of each selected gene product was calculated using the 2^−ΔΔ*Ct*^ method. Additionally, absolute quantification of qPCR products was performed to compare the copy number of miR-132 transcripts expressed in APCs and BM-MSCs. Ten-fold serial dilutions of miR-132 template were performed for the standard curve. Slope and correlation of the standard curve were - 0.33 and 0.99, respectively. Spike-in Cel-miR-39 was added to cell conditioned media (CCM) and explanted swine aortic valves for normalization and quality control (219610, QIAGEN, Germany). All reactions were performed in technical triplicates. All primers and probes are listed in **Supplementary Table I**. APCs from 5 donors and BM-MSCs from 4 donors were assayed in technical triplicates.

### Protein isolation and western blotting

RIPA buffer (R0278, Merck/Sigma-Aldrich, UK) or NE-PER ™ Nuclear and Cytoplasmic Extraction Kit (78833, Thermo Scientific, UK) were used to isolate total protein or cytoplasmatic/nuclear protein fractions for western blotting, following the manufacturer’s protocols. All lysis buffers were supplemented with inhibitors of proteases (1/100 (v/v)) and phosphatases (1/50 (v/v)) (P8340 and P5726, Merck/Sigma-Aldrich, UK). Protein concentration was quantified using a BCA protein assay (23252, ThermoFisher Scientific, UK). Total protein (5–20 μg) was resolved onto 8–12% SDS-PAGE and blotted onto 0.2 μm pore size PVDF membranes (1620177, Bio-Rad, Hercules, CA, USA). Primary antibodies were incubated overnight at 4°C after blocking membranes for 1h, at RT in 5% fat-free milk or 3% BSA dissolved in TBST buffer (100 mM Tris, 150 mM NaCl, pH 7.5 and 0.05-0.1 % Tween-20). ß-Actin or ß-tubulin were used as a loading control for RIPA and cytoplasmic cell lysates; histone H4 or Laminin A/C for nuclear fractions. The list of primary antibodies and titrations used is shown in **Supplementary Table II**. Densitometric band analysis was performed using ImageJ software (National Institutes of Health, Bethesda, MD, USA; https://imagej.nih.gov/ij/) and Image Lab software (Bio-Rad, UK).

### Assays on conditioned media

CCMs were collected and centrrifuged at 10,000 *g* for 3 min, at 4°C to remove cell debris and supernatants were kept at −80°C until analysis. CCMs were used in ELISA, lactate release and glucose consumption assays, for quantification of miR-132, and to condition cells and valves prior to funcional assays.

***ELISAs*** were performed to quantify protein expression in CCM from at least 4 APC and BM-MSC lines, with assays performed in technical triplicates. Immunoreactive levels were normalized to total protein concentration. The following factors were determined: VEGF (DY293B), ANGPT-1 (DY923), BMP2 (DY355) (all from R&D Systems, Oxford, UK).

***Glucose consumption*** and ***lactate release*** were assessed using Glucose-Glo™ assay and Lactate-Glo™ assay (J6021 and J5021, Promega, UK). CCM from 4 APC and BM-MSC lines were assayed in technical triplicates. Appropiate standards and basal media were assayed in parallel. Glucose consumption was calculated as follows: Glucose consumption (mM)_sample_ = [Glucose]_basal media_ - [Glucose]_CCM sample_. Relative Luminiscence Units (RLU) were recorded using a GloMax^®^ Discover Microplate Reader (Promega).

### Cell viability and proliferation in bovine pericardium

APCs seeded on FDA-approved bovine pericardium (BP) and APCs seeded on plastic (2D control) were assessed using fluorescent calcein AM/ethidium homodimer III (EtDHIII) Live/Dead assay (30002-T, Biotium Inc, Insight Biotech, UK) and Click-iT^®^ EdU (5-ethynyl-2’deoxyuridine) Assay (BCK-EDU488, baseclick GmbH, Merck/Sigma-Aldrich, UK), respectively, following the manufacturer’s protocols. Three days before studying proliferation rate, EdU was incorporated to fresh media at 10μM final concentration. Nuclei were counterstained with 300nM 4’,6-diamidino-2-phenylindole dilactate (DAPI) (D1306, ThermoFisher Scientific, UK) for imaging assessment (Zeiss Axio observer Z1 microscope). Images from at least 5 random fields were snapped unless otherwise indicated. In addition, MTS assay (G3582, CellTiter 96^®^ Aqueous One Solution Cell Proliferation Assay, Promega, UK) was performed for colorimetric quantification of viability. All assays were performed on APCs from at least 4 donors in technical quintuplicates.

### 3’-UTR luciferase constructs preparation, molecular cloning, and luciferase assays

Online tools TargetScan, PITA and miRWalk2.0 were used to predict *in silico* the putative 3’UTR target binding sites of miR-132 (hsa-miR-132-3p, MIMAT0000426). UCSC Genome Browser was consulted to obtain the 3’UTR oligonucleotide sequences listed in **Supplementary Table III**. *PmeI* (GTTT/AAAC) and *NotI* (GC/GGCCGC), and *Sa1I* (G/TCGAC) endonuclease restriction sites were inserted into the predicted sequences at 5’ and 3’ ends, respectively, to confirm oligonucleotide clonning into pmiRGLO Dual Luciferase miRNA Target Expression Vector (E1330, Promega, UK). Heat shock transformation was used to amplify luciferase plasmid construct in JM109 *E. coli*. Luciferase plasmid construct preparations were then co-transfected with scramble sequences or miR-132 mimic/inhibitor into 80% confluent HEK293 cell monolayers. Lipofectamin LTX (15338030, Thermo Fisher, UK) was used as lipotransfectant reagent. Luciferase assays were performed using a Dual-Glo^®^ Luciferase Assay System (E2920, Promega, UK) following the manufacturer’s instructions.

### AgomiR and antagomiR assays

APCs were transfected with 25nM antagomiR-132 or agomiR-132 or scramble (Scr) controls using lipofectamin RNAiMAX (13778075, Invitrogen, Thermo Fisher, UK).

### *Ex vivo* model of swine aortic valve calcification

*Ex vivo* valve calcification was modelled in explanted aortic valves (EAV) from 6-month old male pigs using 3mM HP stimulation for 5 or 7 days, as previously described.^24, 25^ Histological assessment and RNA isolation was carried out in harvested sampples. Alizarin Red (ARS), Elastic van Giesson (EVG), Movat or Alcian Blue/Sirius Red (AB/SR) stainings were performed on valves from 4 swine (in duplicate) to quantify calcification, elastin and collagen/proteoglycans. Active osteogenesis was studied by qPCR.

### APC engineering of decellularised bovine pericardium

Commercially FDA-approved glutheraldehide cross-linked BP clinically certified for clinical use was seeded with APCs at increasing seeding density up to 32,000 cell/cm^2^. Unseeded pericardium was used as a negative control. Endpoints were antigenic profile preservation of seeded cells, viability, and proliferation. CCMs were also collected to perform the AorEC scratch ‘wound healing’ assay.. In addition, immunocytochemistry (ICC) analysis was applied to verify preservation of antigenic profile of APCs-engineered on BP

### Statistical analysis

Continuous variables are shown as mean ± standard error of the mean (SEM) or median (IQR) depending on their distribution. Categorical variables are presented as percentages. Normally distributed variables were analyzed using the Student’s t test (two group comparison) or one-way analysis of variance (multiple comparisons ANOVA), as appropriate. Homoscedasticity was assessed with the Levene test, and ANOVA post-hoc analysis included Tuckey or T3 Dunnet testing, as appropriate. Non-parametric tests, including the Mann–Whitney U test or the Kruskal-Wallis test, were used for data not normally distributed. Statistical significance was accepted at p < 0.05. Analyses were performed using SPSS 19.0 for Windows (SPSS, Inc., Chicago, IL, USA) or GraphPad Prism 5.0 (GraphPad, La Jolla, CA, USA)) statistical packages.

## Results

### Antigenic profile of human APCs

Flow cytometry analysis showed that culture-expanded APCs expressed pericyte and mesenchymal markers, such as CD105, CD90, CD44, NG2/CSPG4, and PDGFRβ, while being negative for CD31, CD146, and CD45 (**Supplementary Figure 1A**), in line with previous publications.^15, 26^

### Human APCs are resistant to calcification induced by high inorganic phosphate

During the development of VHD, inorganic phosphate accumulation triggers the transformation of cardiac valves into heterotopic bone-like tissue.^27^ Here, we investigated if high levels of inorganic phosphate in the culture medium could induce osteochondrogenic transition of human APCs. BM-MSCs, used here as an alternative cell comparison, showed accumulation of calcium (Alizarin Red staining) and phosphate (von Kossa staining) after 4 day stimulation with HP, whereas no staining could be appreciated in either assays on APCs after 12 days of HP (**Figure 1A**). A large increase in calcium content was confirmed in BM-MSCs using the *o*-cresoftalein colorimetric method (**Figure 1B**) while negligible readings were reported in APCs (data not shown). Calcification was further evaluated in APCs using a more potent stimulus consisting of 4- and 10-days incubation in DMEM supplemented with 4.5g/mL glucose (25mM) (high glucose, HG) and HP. This combination induced a massive increase in mineralization and cell death in BM-MSCs after only 24h (**Supplementary Figure 1B**). In contrast, calcium deposits were modestly detected in APCs after 10 days, as assessed by Alizarin Red staining (**Figure 1C**) or the *o*-cresophtalein method (**Figure 1D**). Moreover, several molecular readouts of osteoblast differentiation were studied at the mRNA and protein level. QPCR demonstrated that HP induced the acquisition of an osteoblast-like phenotype by BM-MSCs, as indicated by the increase in *BMP2, RUNX2, SOX9, SP7/OSX*, and *SPP1* mRNA expression compared with unstimulated BM-MSCs; whereas, the reverse was seen in APCs, which exhibited downregulated *RUNX2, SOX9, SP7/OSX*, and *SPP1* expression following HP stimulation as compared with the unstimulated condition (**Figure 1E**). Western blotting analysis showed that BMP2 (a morphogenetic protein implicated in bone and cartilage formation), RUNX2 (Runt-related transcription factor 2, a key transcription factor associated with osteoblast differentiation) and OCN (Osteocalcin, a secreted osteoblast morphogen that influences matrix mineralization) were all induced by HP in BM-MSCs, whereas all the studied osteogenic proteins were downregulated in HP-stimulated APCs (**Figure 1F&G**). Moreover, HP increased the levels of BMP2 in BM-MSC-derived CCM. APCs showed secreted BMP2 concentrations comparable to those of BM-MSCs under unstimulated conditions, but they did not secrete extra protein under HP stimulation (**Figure 1 H-I**). A time course of osteoblast marker expression was additionally studied in HP-stimulated APCs, with results showing a general downregulation following extended stimulation (**Supplementary Figure 2A&B**). Moreover, no calcification was detectable at any time point and APC viability was preserved (**Supplementary Figure 2C&D**).

**Figure 1:**
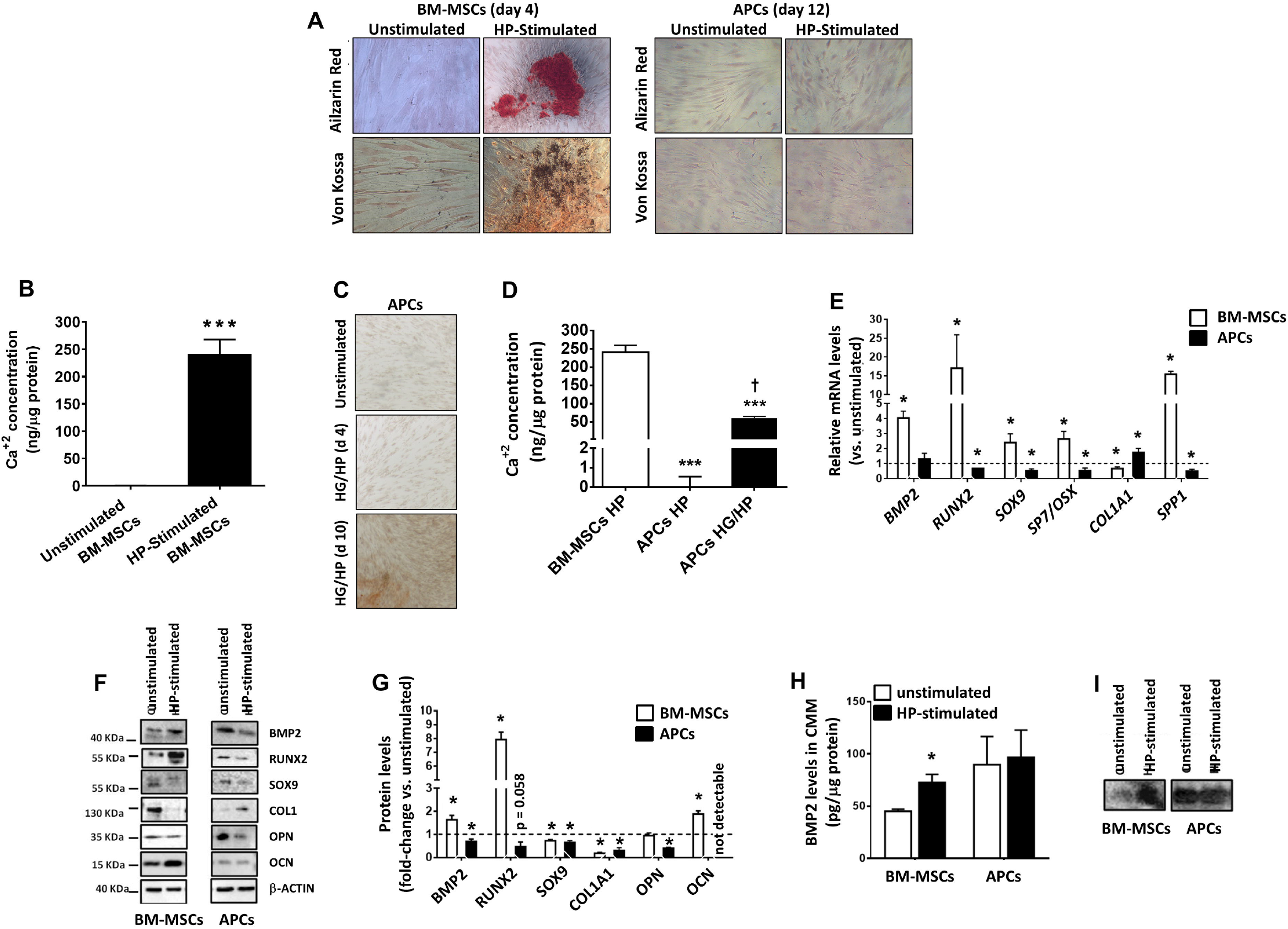
Effect of high phosphate on calcification and osteoblast differentiation. **A**, Representative microscopy photographs of Alizarin Red (calcium deposits) and von Kossa (phosphate deposits) stainings captured from unstimulated and stimulated BM-MSCs (4 days incubation in high phosphate, HP) and APCs (12 days in HP). **B**, Colorimetric quantification of calcium in BM-MSCs normalized by total protein content. ***p<0.001 *vs*. unstimulated. **C&D**, Representative images (Alizarin Red staining) (**C**) and bar graph (**D**) showing the calcium content (assessed using the *o*-cresoftalein assay) in APCs after 5 and 10 days of incubation in a calcifying medium consisting of combination of high glucose (HG) and HP, in comparison with BM-MSCs or APCs stimulated with HP only. ***p<0.001 *vs*. BM-MSCs.^†^p<0.05 *vs*. APCs HP. **E**, Changes in mRNA expression levels of typical osteoblast markers in BM-MSCs and APCs following HP conditioning relative to unstimulated condition in corresponding cell type. *p<0.05 *vs*. unstimulated. **F&G**, Representative western blotting images and bar graph illustrating the results from band densitometry analysis. Data indicate the fold changes in intracellular protein levels following HP conditioning relative to unstimulated condition in corresponding cell type. Band densitometries were normalized by β-actin. *p<0.05 *vs*. unstimulated. **H&I**, Levels of BMP2 in conditioned media (CCM) collected from BM-MSCs and APCs under basal and HP conditions. **H**, BMP2 was quantified in CCMs using ELISA and normalized for total protein content. **I**, Representative image of Western blotting, which shows data in line with the ELISA results. All experiments were performed in cells isolated from 4 different donors, using three technical triplicates. Data are presented as mean ± SEM.

### HP induces miR-132 expression in human APCs

We next investigated if HP regulates the expression of miR-132; the rationale being this microRNA is reportedly relevant for human APC ability to promote tissue repair through molecular mechanisms that can also control extracellular matrix remodelling.^16^ Interestingly, HP induced an early (day 4) up-regulation of intracellular miR-132 (**Figure 2A**) which was sustained until day 12 (**Supplementary Figure 3**). However, miR-132 levels in APC-derived CCM remained unchanged between unstimulated and HP-stimulated conditions (**Figure 2B**). In contrast, BM-MSCs showed reduced intracellular levels and increased extracellular levels of miR-132 in response to HP as compared with the unstimulated condition (**Figure 2C&D**). An absolute quantification of miR-132 copy numbers revealed a large variability regarding BM-MSCs, whereas APCs showed a significant increase in intracellular miR-132 following HP stimulation (**Figure 2E**).

**Figure 2:**
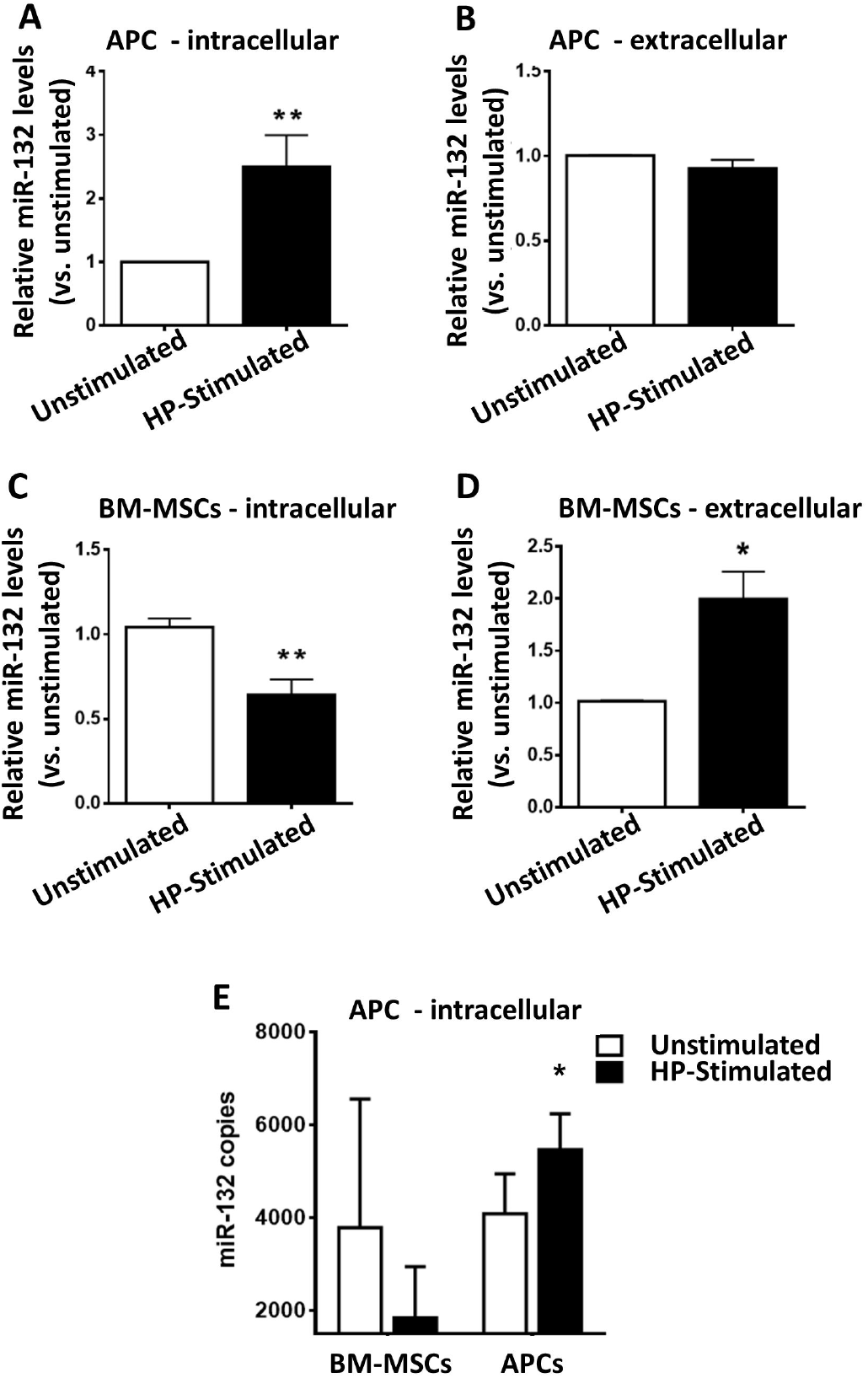
Effect of high phosphate on miR-132 expression. **A**, Intracellular expression levels of miR-132 in APCs under basal conditions (unstimulated) and after 4-day conditioning with high phosphate (HP-stimulated). **B**, MiR-132 content in CCM derived from APCs under basal conditions and after 4-day conditioning with HP. **C**, Intracellular expression levels of miR-132 in BM-MSCs unestimulated and HP-stimulated for 4 days. **D**, MiR-132 content in BM-MSC-derived CCM. **E**, MiR-132 number of copies expressed by APCs and BM-MSCs under unstimulated or HP-stimulated conditions. Intracellular expression of miR-132 was normalized by U6 snRNA expression. Cel-miR-39 spike-in was used during RNA extraction for subsequent normalization in RT-qPCR assays. All experiments were performed in cells isolated from 4 different donors in technical triplicates. *p < 0.05 and **p<0.01 *vs*. basal unstimulated conditions.

Using three bioinformatics tools (TargetScan, PITA, and miRWalk2.0), we found that miR-132 directly targets several genes associated with physiologic and ectopic osteogenic processes, including *CALU, GDF5, ACVR1, GLUT1, MECP2, METLL25, EP300, HBEGF*, and *SMAD7*. Interestingly, qPCR data indicated that HP-induced upregulation of miR-132 expression by APCs was associated with downregulation of all the above candidate targets, apart from *HBEGF* (**Figure 3A**). Luciferase constructs were prepared to validate the regulatory effect of miR-132 on the *in silico* predicted sites (**Supplementary Figure 4A**). Different endonuclease restriction enzyme digestions were used in parallel to validate the insertion of miR-132 targets (**Supplementary Figure 4B**). Luciferase assays were then performed in HEK293 cells co-transfected with seven constructs from the list of predicted genes together with agomiR-132, antagomiR-132, or Scr sequences. Forced expression of the microRNA by agomiR-132 resulted in the downregulation of *CALU, GDF5, ACVR1, GLUT1, MECP2, METTL25*, and *EP300*, as it would be expected for genes that are under inhibitory control; whereas, antagomiR-132 caused a more subtle response, with only three *GDF5, ACVR1* and *EP300* being upregulated following miR-132 inhibition compared with Scr (**Figure 3B-H**). Moreover, after confirming that antagomiR-132 (**Figure 3I**) and agomiR-132 (**Supplementary Figure 5A**) oppositely modulate miR-132 expression in APCs, we asked if miR-132 inhibition could result in expressional changes of the osteogenic factors in APCs. As shown in **Figure 3J**, antagomiR-132 caused a remarkable upregulation of *GLUT1* and induced a significant but milder increase in *CALU* expression under unstimulated conditions. In HP-stimulated APCs, antagomiR-132 induced *CALU, GDF5, AVCR1, GLUT1, MECP2, METLL25* and *EP300*, but not *HBEGF* or *SMAD7* (**Figure 3K**). Following transfection of APCs with agomiR-132, we found that three targets were consistently downregulated both under unstimulated or HP-stimulated conditions (*CALU, GDF5*, and *ACVR1*), two were reduced only under unstimulated conditions (*GLUT1* and *SMAD7*), or under stimulated conditions (*METTL25* and *EP300)*, while no significant effect was seen regarding *MECP2and HBEGF* (**Supplementary Figure 5B&C**).

**Figure 3:**
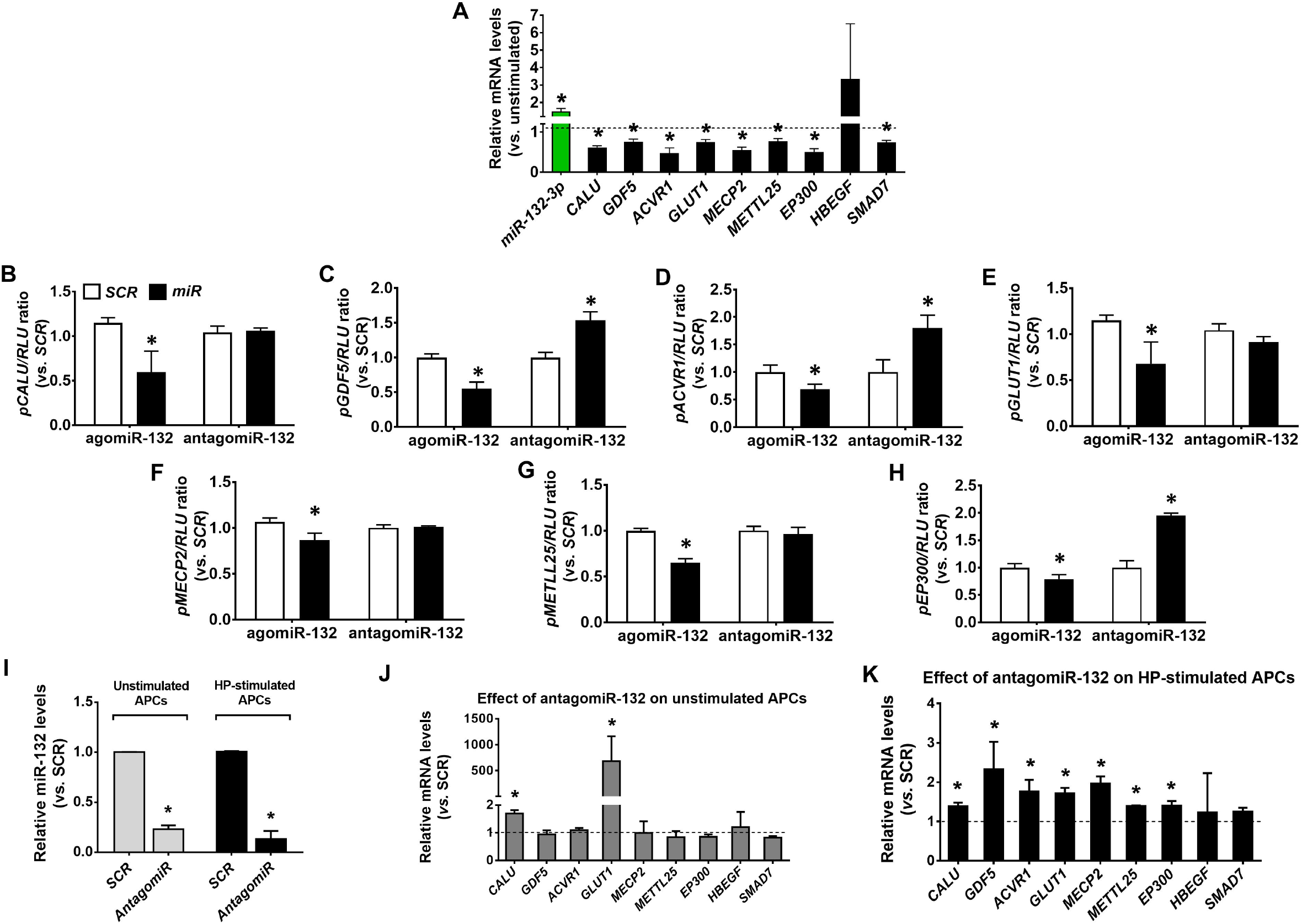
miR132 modulates osteoblastic gene expression in human APCs. **A**, Bar graph showing the expression of candidate target genes in HP-stimulated APCs relative to the expression of unstimulated condition (dotted line). Intracellular levels of miR-132 (miR-132-3p, green bar) were normalized by U6 snRNA expression, while mRNA targets (black bars) were normalized by GAPDH. *p<0.05 unstimulated. **B-H**, Bar graphs showing the results of luciferase assays in HEK293 cells cotransfected with miR-132 mimic (agomiR), miR-132 inhibitor (antagomiR), or Scramble (SCR). RLU for *Renilla sp* luciferase was used as internal control for each reading after 48h of the transfection. *p<0.05 vs. SCR. **I**, miR-132 expression was transiently knocked-down in antagomiR experiments either under unstimulated conditions or following stimulation with HP for 4 days. *p<0.05 vs. SCR. **J&K**, Bar graph showing the effect of miR-132 inhibition on predicted targets under unstimulatedconditions (**J**) and following stimulation with HP (**K**) *p<0.05 vs. SCR. All experiments were performed in APCs isolated from 4 different donors using technical triplicates. Luciferase assays were performed in technical quintuplicates. Data are pepresented as mean ± SEM;

### HP stimulation induces glycolysis in BM-MSCs but not in APCs

Induction of GLUT1 marks the glycolytic switch of stromal cells during osteoblast differentiation.^28^ As shown above, the opposite was seen in APCs, which manifested a reduction in *GLUT1* expression following the HP challenge. This data would suggest there was no glycolytic activation in stimulated APCs. To confirm this possibility, we next investigated the metabolic changes occurring in APCs and control BM-MSCs during forced osteochondrogenic differentiation. In BM-MSCs, HP stimulation increased glucose consumption and lactate and VEGFA secretion, without altering ANGPT-1 release. In contrast, in APCs, glucose and lactate did not change in response to HP (**Supplementary Figure 6A-C**). Moreover, APCs showed lower VEFG and higher ANGPT-1 secreted levels compared with BM-MSCs (**Supplementary Figure 6C&D**).

### Inhibition of miR-132 triggers molecular changes instigating osteochondrogenic transition of human APCs

We next assessed if inhibition of miR-132 could weaken the capacity of APCs to resist forced calcification by HP/HG stimulation. Alizarin red staining revealed mildly increased calcification in miR-132 knocked-down APCs compared with Src-transfected APCs (**Figure 4A**); colorimetric analysis confirmed the inductive effect of miR-132 inhibition on calcium deposits after 5 days exposure to HP/HG (**Figure 4B**). Such an induction resulted in significantly increased glucose consumption and lactate release in miR-132 knocked-down APCs (**Figure 4C&D**). Acquisition of an osteogenic-like profile following miR-132 inhibition was confirmed by the concomitant upregulation of BMP2 and RUNX2 proteins, in association with enhanced GLUT1 expression (**Figure 4 E&F**).

**Figure 4.**
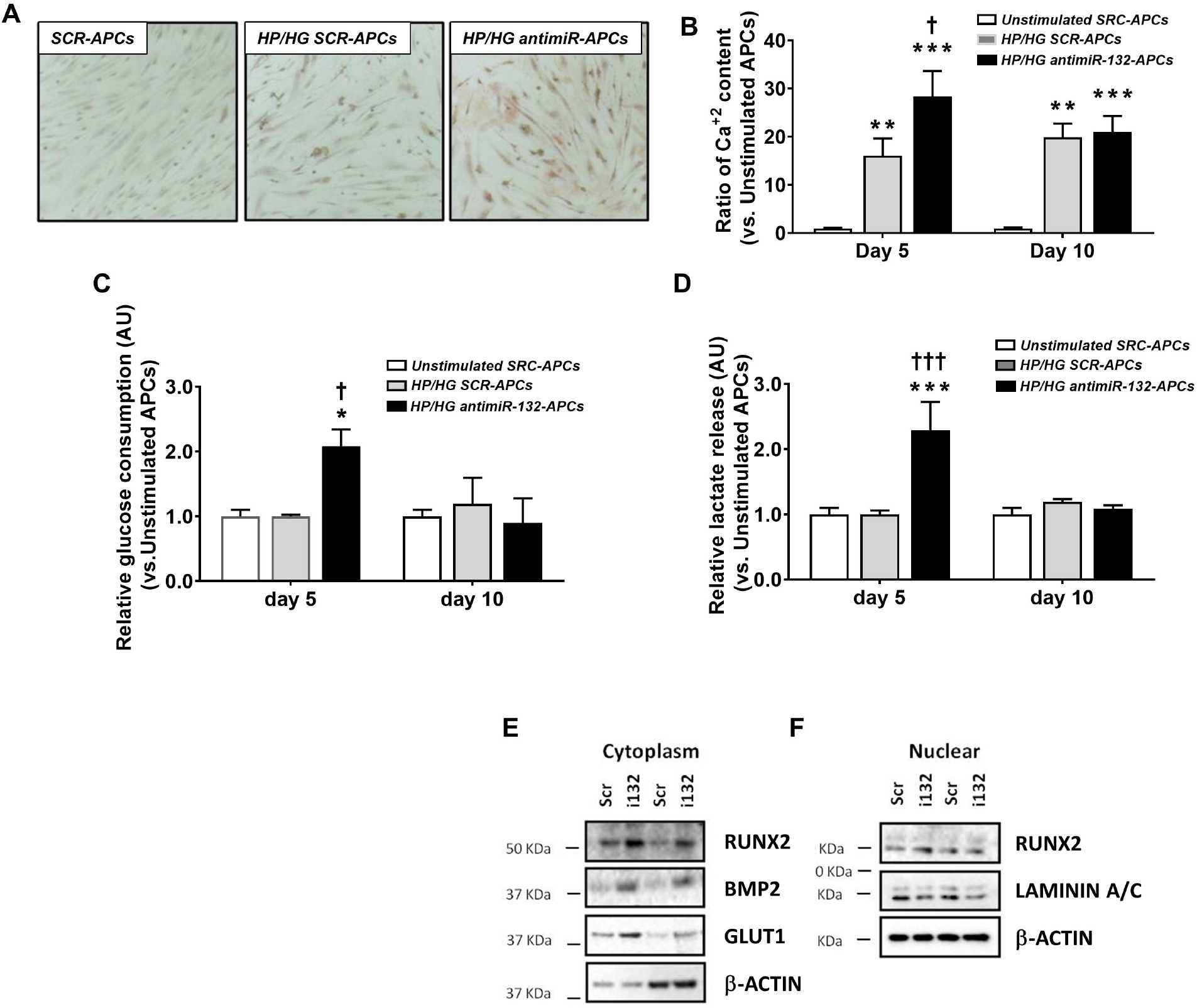
MiR-132 inhibition blunts APC resistance to high phospate/high glucose induced osteogenesis. **A**, Representative microphotographs of Alizarin red stained APCs. Cells were cultured in HP/HG for 10 days, following transfection with antagomiR-132 or scramble (SCR) sequence and compared with unstimulated SCR-transfected APCs. **B**, Colorimetric quantification of calcium deposits in APCs subjected to the same protocol as in (**A**) with measurements performed at 5 and 10 days. **C&D**, Relative glucose consumption and lactate release in APCs subjected to the same protocol as in (**A**) with measurents performed at 5 and 10 days. **E&F**, representative Western blotting of calcyfing markers in APC cytoplasm (**E**) and nuclear (**F**) fractions from 2 different donors. All the other experiments were performed in cells isolated from 4 different donors in technical triplicates. Data are pepresented as mean ± SEM; *p < 0.05, **p<0.01, and ***p<0.001 *vs*. Unstimulated APCs; ^†^p<0.05, and ^†††^p<0.001 vs. HP-Stimulated SCR-APCs.

### The APC-derived secretome prevents osteogenic differentiation of swine aortic valves through miR-132 signalling

Next, we asked if factors secreted by APCs could pass the anti-calcific phenotype to valvular tissue. To this purpose, we tested the effect of the CCM derived from naïve APCs or antagomiR-132 transfected APCs on explanted EAVs exposed for 7 days to HP stimulation. EBM2 and Scr-transfected APC-derived CCM were used as controls (experimental protocol illustrated in **Figure 5A**). EAVs conditioned with naïve APC-derived CCM expressed greater amounts of miR-132 compared with EAVs exposed to EBM2 (**Figure 5B**). In addition, HP stimulation increased the expression of *BMP2, RUNX2, SOX9*, and *SPP1* in EAVs (**Supplementary Figure 7A&B**). The inductive effect of HP on osteoblastic markers was inhibited by the APC-derived CCM (**Figure 5C-F**). AntagomiR-132 transfection reduced the levels of miR-132 in the APC-derived CCM by 65% compared with Scr-transfected APCs (p<0.01, data not shown). Moreover, miR-132 inhibition abolished the ability of APC-derived CCM to induce the expression of miR-132 in EAVs, as demonstrated by a reduction relative to the Scr-transfected APC CCM (**Figure 5G**). Likewise, miR-132 inhibition increased the expression of osteoblastic markers compared with Scr control, apart SOX9 which remained unaltered (**Figure 5H-K**).

**Figure 5.**
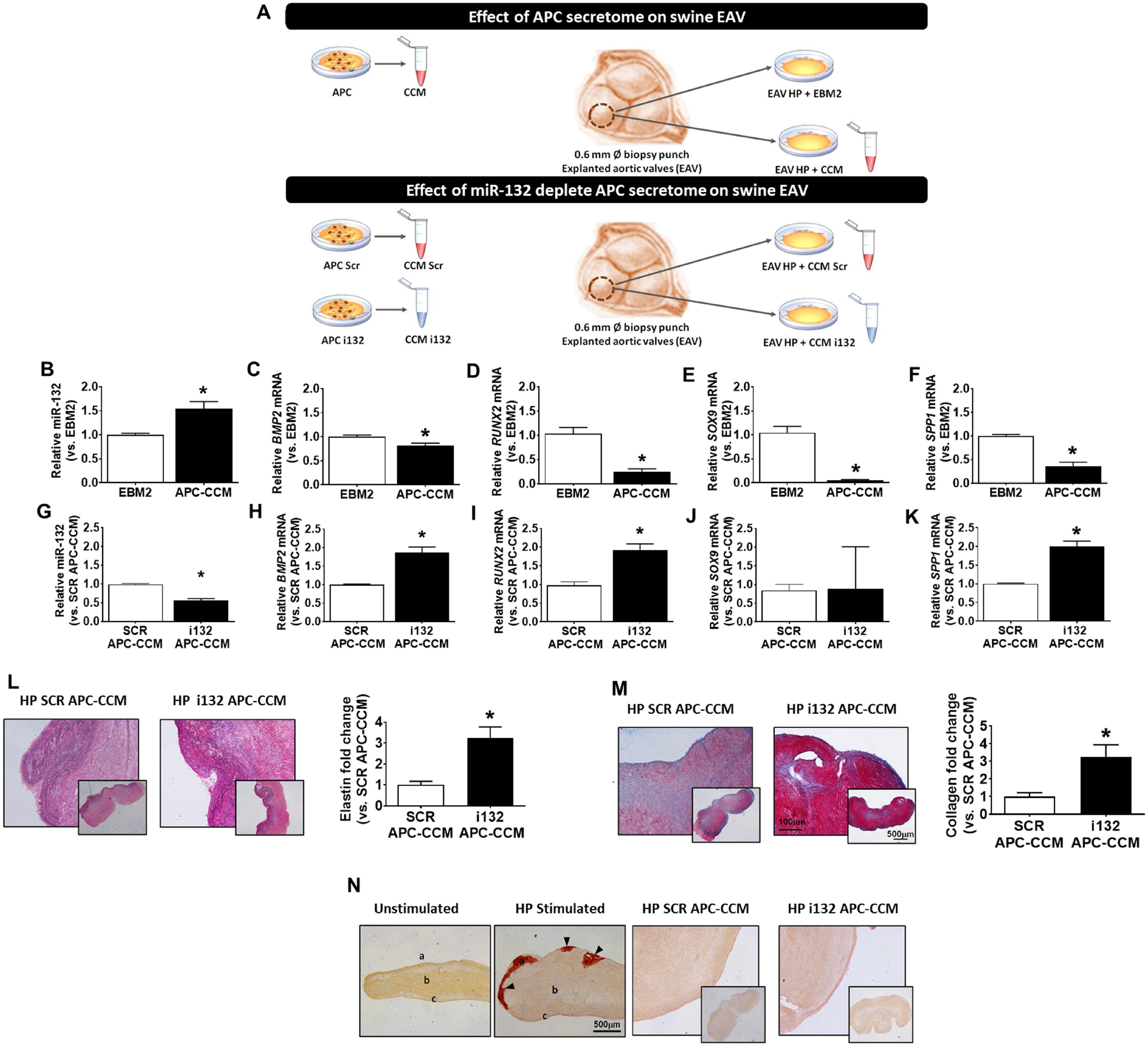
Effect of APC-derived secretome on an *ex vivo* swine model of aortic valve calcification. **A**, Experimental protocol used to assess the effect of APC-derived conditioned medium (CCM) on the explanted aortic valve (EAV) model. In a first set of experiments, the CCM collected from naïve APC was added to the EAV assay. In a second set, APCs were transfected with antagomiR-132 or Scramble (SCR) and then the corresponding CCM (SCR APC-CCM or i132APC-CCM) was added to the EAV assay. In both conditions, EAVs were stimulated with HP (3.2mM), which resulted in induction of osteoblastic markers (see Supplementary Figure 7). **B-F**, Effect of the naïve APC-derived CCM on the relative expression of miR-132 (**B**), *BMP2* (**C**), *RUNX2* (**D**), *SOX9* (**E**) and *SPP1* (**F**) by HP-stimulated EAVs. EAVs treated with EBM2 instead of APC-CCM were used as a control. **G-K**, Effect of the CCM from APCs transfected with SCR or miR-132 inhibitor (i132) on gene expression by EAVs. AntagomiR-132 reduced the miR-132 levels to 0.35-fold the values of SCR-transfected APCs (data not shown). Changes in the expression (relative to SCR APC-CCM) of miR-132 (**G**), *BMP2* (**H**), *RUNX2* (**I**), *SOX9* (**J**) and *SPP1* (**K**) in HP-stimulated EAVs. **L**, Representative images of EVG-stained EAVs stimulated with HP (3.2mM) and treated with SCR APC-CCM or antagomiR-132 APC CCM. Bar graph showing quantitative values. **M**, Alcian blue/Sirius Red staining of EAVs stimulated with HP (3.2mM) and treated with SCR APC-CCM or antagomiR-132 APC CCM. Bar graph showing quantitative values. **N**, Representative microphotographs of Alizarin Red staining showing calcification of EAV exposed to HP as compared with unstimulated condition, and the effect of CCM from SCR or antagomiR-132 transfected APCs. All expreiments were performed in 2 aortic valve biopsies from 4 animals. APC-derived CCMs were pooled to condition each biopsy. Data are pepresented as mean ± SEM; *p < 0.05 *vs*. respective control in each panel.

HP stimulation enhanced elastin (EVG staining, dark purple blue) and collagen content (Movat staining, yellow) in EAVs (**Supplementary Figure 7C**). The above effects were blunted by the naïve APC-CCM, but this action was reversed by miR-132 silencing of APCs (**Figure 5L&M**). Calcification was induced in EAVs following 7 days of HP stimulation, as assessed using the Alizarin Red staining. This effect was remarkably attenuated by the naïve APC-derived CCM as well as by the CCM derived by antagomiR-132 transfected APCs (**Figure 5N**), suggesting the participation of additional factors contained in the CCM.

### Effect of HP on extracellular matrix proteins production by human APCs

Having demonstrated that factors secreted by APCs could inhibit osteochondrogenic transformation of EAVs, we sought to investigate the influence of HP on APC production of collagen and GAG proteins. There was a 1.6-fold increase in collagen protein released by APCs following HP stimulation (**Figure 6A**). APCs are also a natural source of GAGs, mainly N-sulphated, but HP did not produce any significant upregulation of GAGs (**Figure 6 B**).

**Figure 6.**
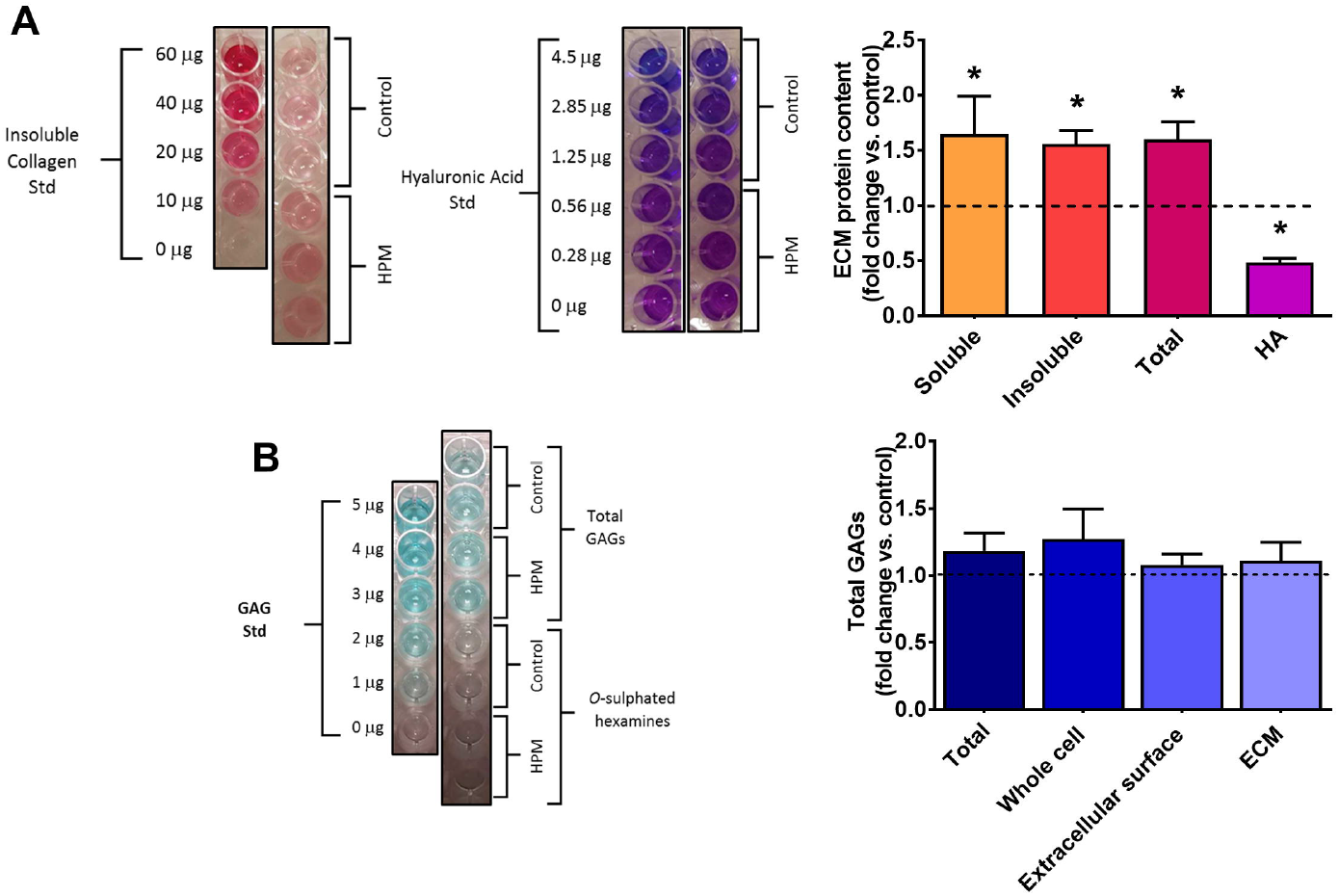
Effect of HP on extracellular matrix proteins production by APCs. **A,** Collagen and hyaluronan (HA) species synthesis in unstimulated (control) and HP-stimulated APCs. Representative images of colorimetric analysis and bar graph showing changes vs. control. **B,** GAGs synthesis in unstimulated (control) and HP-stimulated APCs. Representative images and data of colorimetric analysis. All expreiments were performed in APCs isolated from 3 donors in sixtuplicates. Data are pepresented as mean ± SEM; *p < 0.05.

### Incorporation of APCs into bovine pericardium

Next, we investigated if APCs could be successfully seeded onto FDA-approved decellularized BP. Three different cell densities were tested using APCs from 5 donors, with BP patches being examined after 5 and 10 days of culture in static conditions. The seeding protocol of this and subsequent studies are summarised in **Figure 7A**. APC proliferation (EdU and MTS) and viability (Calcein AM/EtDHIII) in the BP were compared with data obtained from the 2D culture system. At a seeding density of 32,000 cell/cm^2^, BP-seeded APCs showed an excellent viability rate, with constant increase in proliferation rate as compared with day 1 (2.89±0.26 and 1.84±0.29 fold at day 6 and 12, respectively) (**Figure 7B-D**). IHC analysis confirmed the maintenance of the typical antigenic profile by seeded APCs (**Figure 7E**) and their localization within the BP structure (**Figure 7F**). Moreover, in a scratch assay of AorECs performed under HP conditions, the CCM from APC-embedded BP increased the wound closure as compared with the CCM of unseeded pericardium (**Figure 7G**).

**Figure 7.**
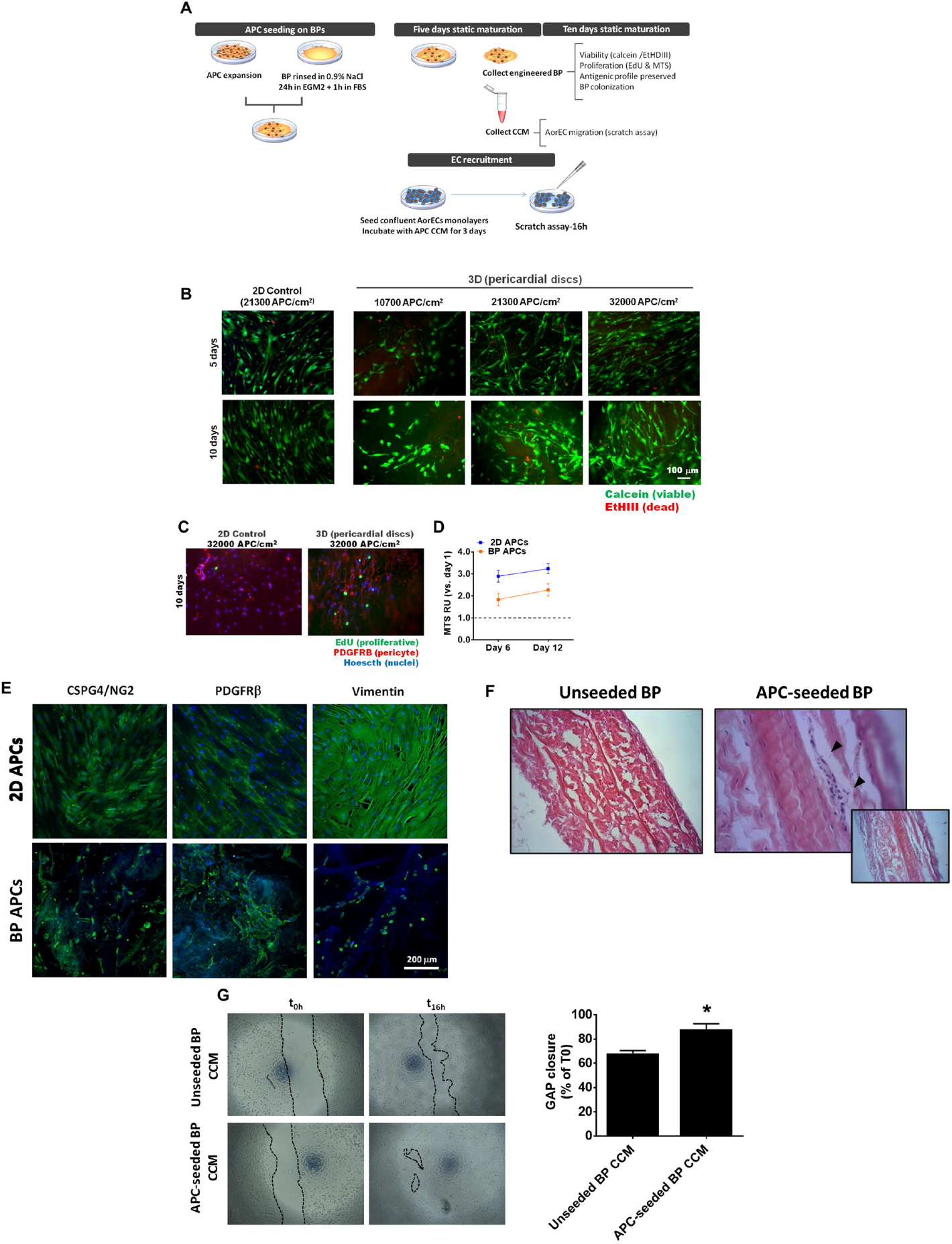
Incorporation of APC on FDA-approved bovine pericardium. **A,** Schematic layout of the experimental protocol. **B,** Representative fluorescence microphotographs of FDA-approved bovine pericardium (BP) seeded with APCs at different densities; images captured at 5 and 10 days from seeding. Living cells are stained green by Calcein AM, while dead cells are stained red by Ethidium Homodimer-III (EtHDIII).**C,** Representative fluorescence microphotographs of EdU-positive proliferating APCs seeded in 2D plates or 3D BP. **D,** plotted MTS reading expressed as changes vs. day 1 values for the 2D and 3D conditions. **E,** Representative fluorescence microphotopgraphy showing antigenic profile retention by APC embedded on BP. **F,** Representative hematoxylin/eosin image of APCs seeded on BP. **G,** Assessement of the chemotactic properties of CCM from APC-embedded BP on AorECs in a scratch ‘wound healing’ assay. Representative image and plotted % GAP closure compared with time 0 (T0). Data are pepresented as mean ± SEM; *p<0.05.

## Discussion

This is the first study reporting the use of human APCs to engineer an FDA-approved BP patches used to manufacture biological prosthetic heart valves; the main findings being (1) the superior ability APCs to resist to osteochondrogenic induction by HP through a mechanism involving the miR-132 and (2) the capacity of APCs to transfer the advantageous phenotype to swine cardiac valves. Importantly, APCs were expanded from saphenous vein tissue leftovers of patients undergoing coronary artery bypass surgery. The donors of BM-MSCs, used as a cell comparison, were younger than APC donors and did not have clinical signs of cardiovascular disease. Therefore, the superior profile of APCs cannot be attributed to a healthier condition of the donor subjects. Finally, we showed the feasibility of using pericytes for cellularization of valvular prostheses; results being in line with our previous data on pericyte-engineered CorMatrix grafts for the correction of congenital heart defects.^17^

Autologous, cell-based, tissue-engineered heart valves with regeneration potential have been suggested to overcome the limitations of currently used bioprostheses, in particular their tendency towards calcific degeneration. BM-derived mononuclear cells, which contain a cocktail of progenitor cells and leukocytes, have been incorporated in minimally invasive protocols for both cell harvest (sternal aspirate) and valve delivery (through a mini-sternotomy or transcatheter aortic valve implantation).^9, 29^ More defined cell populations such as MSCs have been also proposed. MSCs can be easily harvested and expanded from multiple tissue sources. Accordingly, they represent the cell of choice, with several studies suggesting the therapeutic utility in animal models.^8, 30, 31^ Comparison studies attempted to determine the superiority of MSCs vs. other cell types, such as BM-MNCs or CD133^+^ aortic-derived cells.^10, 32, 33^ Nonetheless, these assessments focused on functional readouts, but did not pay much attention to the mechanistic rationale for choosing a specific cell population.

Here, we provide multiple levels of evidence for the superior capacity of human APCs to resist osteochondrogenic induction by HP. In contrast to BM-APCs, APCs showed a downregulation of major osteogenic markers following HP challange, including BMP2, RUNX2, SOX9, COL1A1, OPN, and OCN. These factors form an integrated molecular network relevant for the development and progression of VHD.^8^ Recent studies have pinpointed the key role of miRs during the pathogenesis of cardiovascular calcification. Cui et al showed miR-204 acts as a central regulator of vascular smooth cell calcification by targeting Runx2.^34^ MiR-141 reportedly inhibited the osteogenic differentiation of porcine valvular interstitial cells through a BMP2 dependent pathway.^35^ A miR microarray study showed that miR-638 inhibits human aortic valve interstitial cell osteogenic differentiation by inhibiting Sp7 transcription factor.^36^ Previous studies have highlighted the importance of miR-132, a highly conserved miR transcribed from an intergenic region on human chromosome 17, in tumoral angiogenesis^19^, cardiac adverse remodelling following aortic coartation^18^, and benefit of APC therapy in a murine model of myocardial infarction^16^. To the best of our knowledge, this is the first report indicating miR-132 plays a key role in the human APC resistance to calcification. Results from studies of gene expression following HP stimulation, *in silico* analysis, luciferase assays, and silencing/forced expression of miR-132 level indicate that this miR could counteract several downstream targets relevant to epigenetic mechanisms (methylation [MECP2 and METTL25] and acetylation [EP300]), transcriptional regulation of the TGF-(j/SMAD signaling pathway (GDF5 and ACVR1) and metabolism (GLUT1). The full list of these genes and related function is reported in **Table 3**. The inhibitory effect of miR-132 on GLUT1 indicate that APCs may have the capacity to modulate glycolysis in response to HP stimulation. Noteworthy, miR-132 was recently described to mediate the ‘metabolic shift’ of prostate cancer cells *via* GLUT1 regulation.^37^ Moreover, GLUT1 is overexpressed during osteoblast differentiation, which underlies the acquisition of the glycolytic profile necessary for increased energy requirements of such a demanding transformation.^28^ Moreover GLUT1 suppresses the AMPK-dependent proteasomal degradation of RUNX2, prolonging its activation.^28^ In line with these concepts, we report that BM-MSCs, but not APCs, showed increased glucose consumption and lactate release in response to HP. Importatly, inhibition of miR-132 caused an increase of both the glycolytic markers and well as an induction of GLUT1, RUNX2, and BMP2 in HP-stimulated APCs. BMP2 is a strong osteogenic morphogen and an upstream regulator of RUNX2, which in turn acts as a master transcription factor regulating the expression of SPP1, alkaline phosphatase, Osterix/SP7, and BGLAP, thereby promoting osteoblast differentiation. Altogether, these results suggest that the expression of miR-132 by APCs is essential to inhibiting the metabolic switch during osteogenic transition.

**Table 3.**
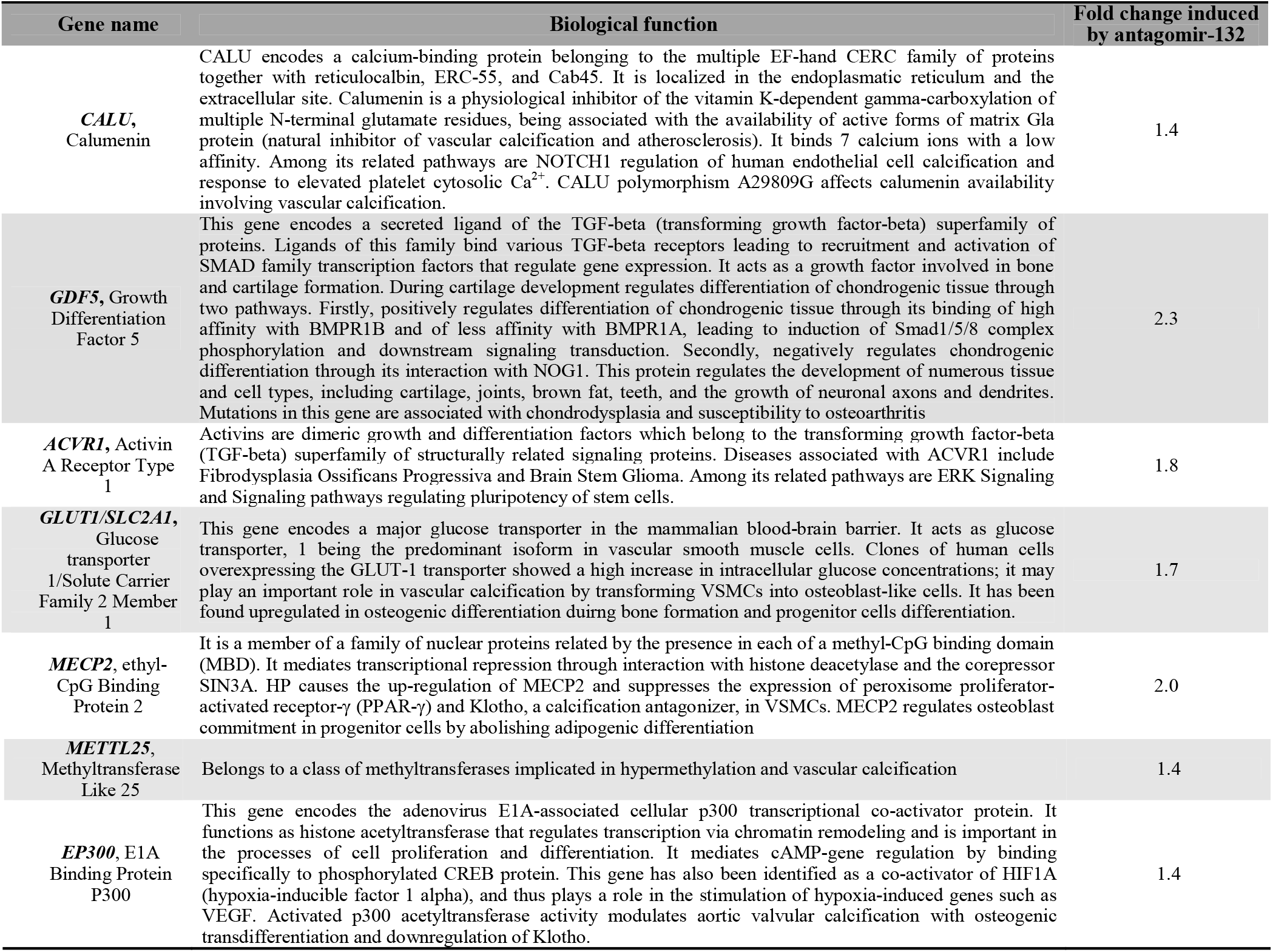
Targets of miR-132 in human APCs

Porcine and bovine pericardial valves are the most commonly implanted prostheses for aortic valve replacement surgery in the UK; and there is hope that immunologic barriers, which contribute to valve degeneration, could be overcome with the use of tissues from genetically modified pigs lacking xenogeneic antigens.^38, 39^ Cellularization with autologous or off-the-shelf allogeneic pericytes could help in this endevour, as evidence points to the immunomodulatory properties of these cells.^40^ In addition, we show for the first time that pericytes from the human vasculature secrete factors that can convey anti-calcific cues to explanted swine valves. The involvement of miR-132 is likely considering that (1) valves exposed to the APC-derived CCM showed an increase in miR-132 levels together with resistance to osteochondrogenic transformation, and (2) antagomiR treatment of APCs precluded these benefits. It should be noted that the inhibition of calcium deposit by the APC-derived CCM was maintained after antagomiR-132 treatment. Therefore, it is possible that other factors contained in the CCM have participated in transferring anticalcific properties to the valve. Nonethless, the antagomiR treatment could not inhibit miR-132 completely; the residual microRNA content could have been enough to inhibit valve calcification. It remains to be elucidated whether the valvular miR-132 was captured from the APC-CCM or was endogenously produced by the swine valve.

Cardiac pericytes have been successfully used to engineer xenogenic material currently employed for cardiovascular reconstructive surgery.^17^ The present study confirms the capacity of vein-derived pericytes to home and proliferate after seeding onto BP patches approved for clinical use and to secrete factors enabling the recruitment of endothelial cells, even in the presence of HP concentrations.

### Perspectives and limitations

Altogether, these findings provide several exciting lines of new evidence pointing towards novel bioengineering solutions based on human pericytes for the treatment of valvular defects. There were limitations in the study. Although the effect size favouring of APCs vs. BM-MSC was large, the number of biological replicates was small, thus requiring replication in larger cohorts. The primary endpoint of the study was to provide a clear and strong mechanistic rationale for the use of pericytes; thereby establishing a solid and ethically acceptable foundation for preclinical studies in large animal models. We are convinced that the evidence collected so far warrants to pursue such an *in vivo* validation as a step toward the clinical use of novel bioengineered prostheses.

## Acknowledgments

Graciela Sala-Newby (PhD) and Thomas Hathway (Mlt) from Bristol Medical School (Translational Health Sciences, University of Bristol) helped with the expansion of pDNA constructs. Hannah Martin (Mlt) and Paul Savage (Mlt) from Bristol Medical School (Translational Health Sciences, University of Bristol) provided technical support for histological sample processing. Patient enrollment and sample collection was performed by research nurses and administrators from the NIHR Biomedical Research Centre at University Hospitals Bristol NHS Foundation Trust and the University of Bristol. Dr. Niall Sullivan from Avon Orthopaedic Centre, Southmead Hospital (Bristol, U.K) collected bone marrow samples.

---

**Supplementary Figure 8.**
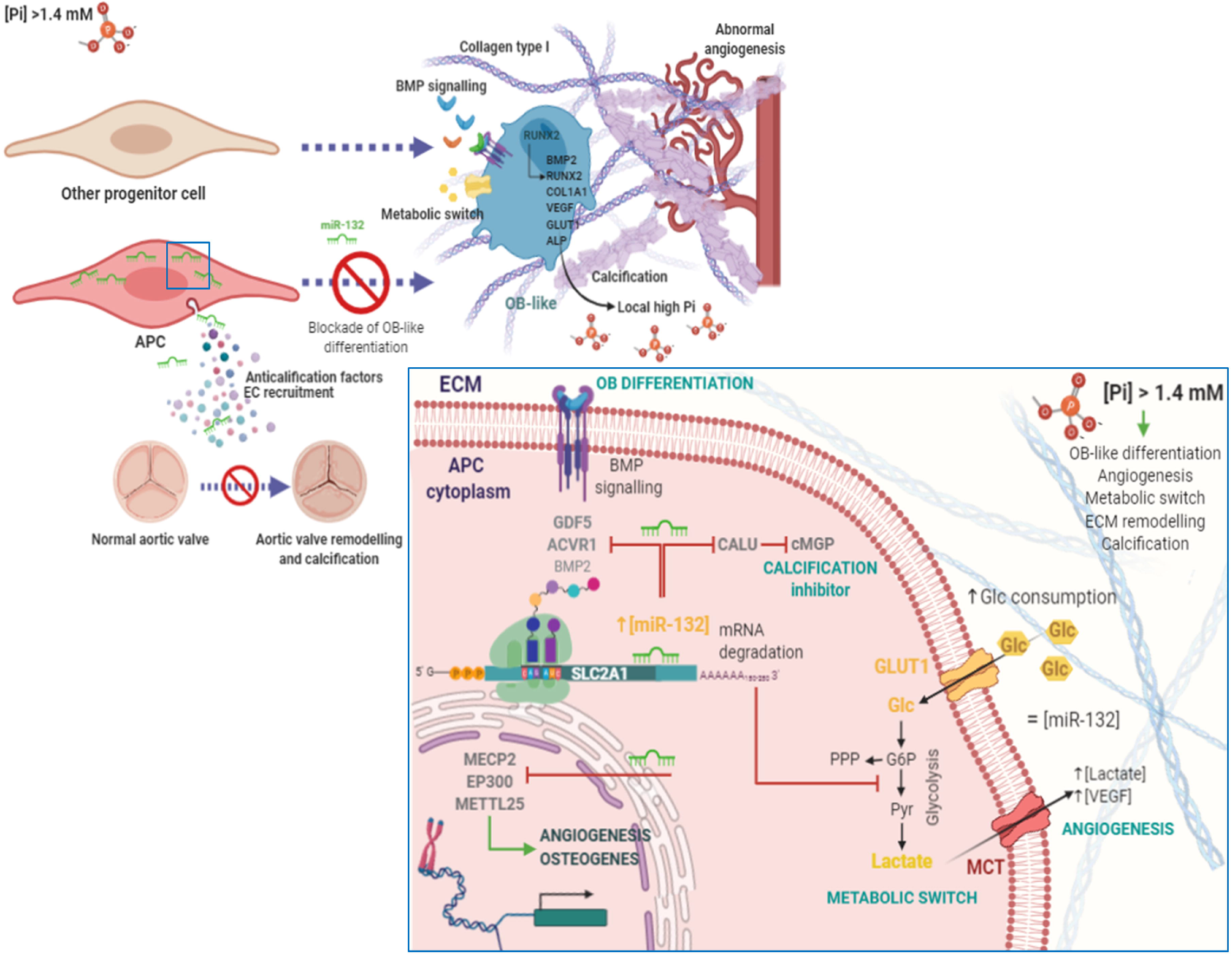

