## supplemental material for "Human saphenous vein provides a unique source of anti-calcific pericytes for prosthetic cardiac valve engineering"

**Methods**

**Clonning methods**

***3′-UTR luciferase constructs and molecular cloning of DNA***

Online tools TargetScan, PITA and miRWalk2.0 were used to *in silico* predict the putative 3’UTR target binding of miR132 (hsa-miR-132-3p, MIMAT0000426). An extensive revision of the literature identified for potential targets in osteogenic differentiation such as CALU, BMP2, GDF5, ACVR1, MECP2, METLL25, EP300, SLC2A1/GLUT1, HB-EGF, SMAD7. UCSC Genome Browser was consulted to obtain the 3’UTR oligonucleotide sequences listed in **Supplementary Table III**. *PmeI* (GTTT/AAAC) and *SalI* (G/TCGAC) endonuclease restriction sites were inserted into the predicted sequences at 5’ and 3’ ends, respectively. In addition, *NotI* (GC/GGCCGC) site was inserted to further confirm oligonucleotide clonning into pmiRGLO Dual Luciferase miRNA Target Expression Vector (E1330, Promega, UK). 3’UTR oligonucleotides were annealed at 40ng/uL (3 μM) each in 1X T4 ligation buffer (B0202S, New England Biosciences, UK), 95 °C for 5min followed by a slow cooling down (5°C/min) to 25 °C. Plasmid preparations (10-20 μg) were double-digested with a total of 2U PmeI-HF and 2U Sal-HF (R0560S and R3138S , New England Biosciences, UK) for 5h, 37°C. Digestion products were then electrophoresed into 1% agarose gel in 1X TBE. Bands were isolated using an Isolate II PCR and Gel kit (BIO-52059, Bioline, UK). Ligation of 285 pg annealed oligos (insert) and 50 ng linealized pmiRGLO was performed in 1X T4 ligase buffer and 1U T4 ligase (EL0013, Thermo Fisher Scientific, UK) for 1h, 20°C to produce Luciferase plasmid constructs. The amount of insert was calculated as follows (ng insert = (molar ratio insert:pDNA x ng pDNA) x (bp insert/bp pDNA)) and the molar ratio chosen was 1:1. Bacterial heat-shock transformation was made in 15 μL JM109 *E. coli* competent cells (L2001, Promega, UK) using 1.5μL of the ligation product. Briefly, transformation was carried out at 42°C for 45 s, followed by 2 min on ice. Transformation product was then 10-fold diluted in SOC media (B9020S, New England Biosciences, UK) and incubated for 60min in agitation before platting into selective LB-Agar (100 μg/mL ampicilin) and further expanded into mini or midi format as appropiate. Plasmid preparations, mini and midi endotoxin free (Merck/Sigma-Aldrich, UK), were assayed to isolate pDNA constructs. Single NotI, and double BahmI + XbaI and BahmI + PmeI (R3189S, R3136S, R0145S, R0560S. All from New England Biosciences, UK) digestions followed by 1% agarose gel electrophoresis were performed to validate insert ligation into pmiRGLO.

***Transient transfection and luciferase assay***

Luciferase plasmid construct preparations were co-transfected with Scr or miR-132 mimic/inhibitor in 80% confluent HEK293 cell monolayers ( 96 multi-well plates). Luciferase plasmid constructs (0.12 μg) were co-diluted with 35 ng (25 nM final concentration) of antagomiR132/antagomiR scramble or agomiR132/agomiR scramble in 5 μL Opti-MEM® I Reduced Serum Medium (31985-070, Gibco^TM^, Thermo Fisher Scientific, UK) and 0.145μL Reagent Plus (1:1 with pDNA/miR diluted). Lipofectamin LTX (0.22 μL) (Invitrogen, Thermo, UK) was also diluted in 5μL Opti-MEM®. Both preparations were incubated for 5 min at 20°C, mixed and incubated for further 15 min. 10μL of the lipocomplexes were inoculated in 100μL final volume and were incubated for 5 h, at 37°C, 5% CO_2_. Luciferase assays were performed after 36 h using a Dual-Glo® Luciferase Assay System (E2920, Promega, UK) following the manufacturer’s instructions. Firefly luminiscence was normalizaed by *Renilla sp*. luminiscence, and ago/antagomiR scrambles were used as 100% RLU for all calculations. Empty pmiRGLO control was additionally transfected with and without ago/antagomiRs to rule out any confounding off-target binding of miR-132 to the backbone. Experiments were performed in three independent experiments. Each experiment was carried out in at least 4 technical replicates per condition. RLU from technical replicates and condition was averaged. The averages were then compared for appropriate statistical analysis.

**AgomiR and antagomiR assays**

APCs were seeded at 10,500 cell/cm^2^ in EGM2 media (12 multi-well plates). After 24 h antagomiR132 (i132) or agomiR132 experiments were conducted by transiently transfecting 25 nM miR132 inhibitor, miR132 mimic or scramble (Scr) controls. Lipofectamin RNAiMAX (13778075, Invitrogen, Thermo, UK) was used following the manufacturer’s protocols. In brief, 3μL of lipofectamin RNAiMAX and miRs (25 nM) were diluted into 50μL Opti-MEM® I Reduced Serum Medium (31985-070, Gibco^TM^, Thermo Fisher Scientific, UK), incubated for 5 min and combined (20 min) to form transfection lipocomplexes. 100μL of the lipocomplexes were inoculated into 500μL final volume and incubated for 6 h, at 37°C, 5% CO_2_. Experiments started after 24h. APCs from 5 donors were usedand were assayed in technical triplicates.

***Ex vivo* model of swine aortic valve calcification**

Aortic valves from untreated 6-month old male pigs were collected into DMEM (4.5g/L glucose) (31966-021, Gibco^TM^, Thermo Fisher, UK) supplemented with 100 U/mL penicillin and 0.1mg/mL Streptomycin (DE17-602E, Lonza, UK). Explanted aortic valves (EAV) were cleaned up and clots removed in PBS. At least 2 biopsies per leaflet were obtained using a 6mm diameter biopsy punch were assayed into 96 multi-well plates for 4 or 8 days. Valve calcification was induced by the addition of 3 mM HP. Harvested samples were destinated either to RNA isolation (at least 2 per valve and condition) and histology (at least 2 per valve and condition) at endpoint. Alizarin Red (ARS), Elastic Verhoeff-Van Giesson (EVG), Movat or Alcian Blue/Sirius Red (AB/SR) stainings were performed on paraffin-embedded EAVs . Quantitative analysis of the extent of calcification, elastin deposition or collagen/proteoglycans was performed using ImageJ software. Leaflets from 3-4 animals were assayed in each set of experiments in technical duplicates.

**APC engineering on decellularised bovine pericardium (APC-BP)**

Commercially FDA-approved glutheraldehide cross-linked bovine pericardium (BP) clinically certified for SVR and valve reconstruction was used to bioengineer APCs on. BP patches were washed in sterile 0.9% saline solution, following the manufacturer’s instructions and were cut into 6 mm diameter discs thereafter using a biopsy punch. BPs consist of one smooth surface and one rough surface. APCs were only seeded on the rough surface and kept in static culture for 5 and 10 days. EGM2 media with no heparine and no VEGF was used, unless otherwise indicated. Three different APC titrations were used: 3,200 cell/disc (10,600 cell/cm^2^), 6,400 cell/disc (21,300 cell/cm^2^) or 9,600 cell/disc (32,000 cell/cm^2^). Unseeded BP discs were used as negative control. BPs were prepared to assess APC function including proliferation, viability, antigenic profile preservation and capability of endothelial cell recruitment in transwell migration assays. CCMs were also collected to perform AorEC scratch ‘wound healing’. Experiments were performed in APCs from 4 donors in duplicate or triplicate.

**Hystological and immunohistochemistry assessment of EAVs and APC-BPs**

EAVs were harvested, fixed in buffered 4% PFA and embeded in paraffin, while APC-BPs were prepared in 1.6% agarose prepared in buffered 10% formalin to avoid cell loss during sample processing prior to paraffinization. Five μm sample sections were depparaffinized to distilled water. Hematoxylin/eosin staining was used in APC-BPs to demonstrate APC migration into the BP. ARS, EVG and Movat pentachrome or AB/SR stainings were used in EAVs. Sample calcification was assessed by staining with 2% (w/v) **Alizarin Red** aqueous solution for 3 min. Alizarin Red solution pH was adjusted to pH 4.1–4.3 with 10% NH_4_OH (A5533; Merck/Sigma-Aldrich, UK). Samples were then dehydrated in acetone, acetone:xylene (1:1) and xylene. Elastin synthesis was proven using **EVG staining**. Fresh Verhoeff’s working solution was prepared with 2.5:1:1 of 5% (w/v) alcholic hemaatoxylin, 10% (w/v) aqueous ferric chloride and Weigert’s iodine solution. Samples were incubated for up to 1 h, rinsed twice in tap water and differentiated in 2% (v/v) aqueous ferric chloride for 2 min. Samples were then incubated with 5% (w/v) sodium thiosulfate for 5 min and counterstained in van Giesson’s solution (1% (v/v) aqueous acid fuchsin in saturated picric acid. ECM composition was assessed using a **Movat pentachrome staining kit** (ab245884, Abcam, UK), following the manufacturer’s instructions. A combination of **Alcian Blue and Sirius Red stainings** was performed to analyse the whole collagen/proteoglycan synthesis in HP EAVs. Alcian blue solution was prepared at 1% (w/v) in 3% acetic acid, pH 2.5 and incubated for 20 min; Sirius Red/Direct Red 80 was prepared at 0.1% in picric acid and incubated for 30 min thereafter. All reagents were provided by Merck/Sigma-Aldrich (UK).

**Immunohistochemistry** techniques were applied in EAVs to validate the regulation of BMP2 in HP samples of EAV. Sodium citrate buffer (10 mM, pH 6.0) was used for heat-induced epitope retrieval (Lab Vision™, Thermo, UK) in a water bath set at 100°C for 15 min and cooled down for 1 h. Antigenic blockade and permeabilization was performed by incubating wih 1% BSA or 5% FBS in 0.1% Tween 20 PBS at 20°C. Primary antibody Bmp2 (1:100, ab230498, Abcam, UK) was incubated for 16 h, 4°C. Anti-rabbit IgG antibody conjugated to Alexa-Fluor 488 (1:400 in PBS) (A32731, Invitrogen, ThermoFisher Scientific; UK) was incubated for 1h at 20°C. Nuclei were counterstained with DAPI (1:1000 in PBS) (D1306, ThermoFisher Scientific, UK) for 3 min.

Slides were mounted using either DPX mounting media (06522, Merck/Sigma-Aldrich, UK) for light mycroscopy approaches or Fluoromount-G®(100241-874, Southern Biotechnology MS, VWR) for immunofluorescent techniques. Imaging was performed using a Zeiss AxioObserver.Z1 fluorescence microscope, unless otherwise specified.

**Extracellular matrix composition**

Confluent APC monolayers were cultured in control or HP conditions into 6 multi-well plates for 8 days to ensure for ECM formation and cross-linking. Pepsin-acid soluble and insoluble collagen fractions were assessed using a Sirius red dye binding assay (Sircol soluble collagen assay-S1000 and Sircol insoluble collagen assay-S2000; Biocolor, UK). In brief, cell monolayers were incubated overnight with 0.1 mg/ml pepsin prepared in 0.2 M acetic acid and centrifuged at >15,000*g* for 10 min. Pellets were kept quantifying insoluble collagen. Supernatants were concentrated overnight at 4°C following the manufacturer’s recommendations to quantify the amount of pepsin acid–soluble collagen. Collagen content was stained with Picrosirius Red solution. Unbound dye from the collagen-dye pellet was removed with acid-salt wash reagent. The remaining pellet yielded a red-to-orange solution after alkali reagent addition, quantified at λ = 550 nm.

Overall PGs and GAGs content were analysed by assaying the content of hyaluronic acid or hyaluronan (HA) and sulphated PGs or GAGs. For HA quantification purposes APC monolayers were harvested with 100μL 5U (50 μg) of proteinase K (P4850, Merck/Sigma-Aldrich, UK) diluted in 50 mM Tris-HCl, pH 7.5 and incubated at 55°C, 16 h. For PG/GAG quantification, APC monolayers were harvested in 1 mL of Papain solution consisting of 0.1 mg/mL (2.6 U/mL) Papain (P3125, Merck/Sigma-Aldrich, UK) dissolved in 0.2 M sodium phosphate buffer pH 6.4, 0.8 % (w/v) sodium acetate, 0.4 % (w/v) EDTA and 61.5 mg/mL Cysteine-HCl. Three aliquots were prepared to assess (1) external cell surface PG/GAGs, (2) total cell content of PG/GAGs, and (3) ECM PG/GAG content. Purple-Jelley Hyaluronan Assay and Blyscan sulfated GAG Assay (Biocolor, UK) were used following the manufacturer’s instructions. *N*-sulfated GAG content was subtracted to total GAGs to calculate the amounts of *O*-sulfated GAGs. ECM composition of HP conditioned APCs was normalised to control APCs and have been shown as percentages (%).

**Supplementary Tables & Figures**

**Supplementary Table I. List of primers and probes used in the manuscript.**

| **Manufacturer** | **Product** | **Application** |
| --- | --- | --- |
| Applied Biosystems™ | U6 snRNA (NR_004394) (assay ID: 001973) | Taqman (Hs, Ss)- HK for miRNA expression |
| Applied Biosystems™ | hsa-miR-132-3p (assay ID 000457 | Taqman (Hs, Ss) |
| Applied Biosystems™ | Cel-miR-39-3p (assay ID 000200 | Taqman- HK for miRNA expression |
| QIAGEN | QT00012544 (Hs_BMP2_1_SG) | SYBR Green (Hs) |
| QIAGEN | QT00020517 (Hs_RUNX2_1_SG) | SYBR Green (Hs, Ss) |
| QIAGEN | QT00001498 (Hs_SOX9_1_SG) | SYBR Green (Hs, Ss) |
| QIAGEN | QT00213514 (Hs_SP7_1_SG) | SYBR Green (Hs) |
| QIAGEN | QT00037793 (Hs_COL1A1_1_SG) | SYBR Green (Hs) |
| QIAGEN | QT01008798 (Hs_SPP1_1_SG) | SYBR Green (Hs) |
| QIAGEN | QT00003087 (Hs_CALU_1_SG) | SYBR Green (Hs) |
| QIAGEN | QT00199864 (Hs_GDF5_1_SG) | SYBR Green (Hs) |
| QIAGEN | QT00071743 (Hs_ACVR1_1_SG) | SYBR Green (Hs) |
| QIAGEN | QT00068957 (Hs_SLC2A1_1_SG) | SYBR Green (Hs) |
| QIAGEN | QT00213514 (Hs_SP7_1_SG) | SYBR Green (Hs) |
| QIAGEN | QT00088214 (Hs_METTL25_1_SG) | SYBR Green (Hs) |
| QIAGEN | QT00094500 (Hs_EP300_1_SG) | SYBR Green (Hs) |
| QIAGEN | QT00000455 (Hs_HBEGF_1_SG) | SYBR Green (Hs) |
| QIAGEN | QT00076391 (Hs_SMAD7_1_SG) | SYBR Green (Hs) |
| QIAGEN | QT00079247 (Hs_GAPDH_1_SG) | SYBR Green (Hs)- HK for mRNA expression |
| QIAGEN | QT00199367 (Hs_RRN18S_1_SG) | SYBR Green (Hs, Ss)-HK for mRNA expression |
| Merck/Sigma-Aldrich | SsBmp2 (Fw: 5'-CTGCGGTCTCCTAAAGGTCG-3'; | SYBR Green (Ss) |
|  | Rv: 5'-AGCAGCAACGCTAGAAGACA-3') |  |
| Merck/Sigma-Aldrich | SsSpp1 (Fw: 5'-CAGACTTTCCTAGCGCCACA-3'; | SYBR Green (Ss) |
|  | Rv: 5'-CTTGCTTGGCAGGGTCTCTT-3') |  |
| Merck/Sigma-Aldrich | SsBglap (Fw: 5'-CTCACACTGCTTGCCCTACT-3'; | SYBR Green (Ss) |
|  | Rv: 5'-CACTAGGCTTTGCATCTGCC-3') |  |

HK, housekeeping gene; Hs, *Homo sapiens*; Ss, *Sus scrofa*

**Supplementary Table II. List of antibodies used in the manuscript.**

| **Manufacturer** | **Product** | **Application (dilution)** |
| --- | --- | --- |
| Merck/Sigma-Aldrich, UK | Ms mAb anti-β-Actin antibody (A5441) | WB (1:30,000) |
| Cell Signalling Technology, UK | Ms mAb anti-β-Tubulin (D3U1W) | WB (1:1,000) |
| Cell Signalling Technology, UK | Ms mAb anti-Lamin A/C (4C11) | WB (1:1,000) |
| Abcam, UK | Rb pAb anti-BMP2 (ab14933) | WB (1:500) |
| Abcam, UK | Rb pAb anti-proBMP2 antibody (ab230498) | 1:500 (WB), 1:100 (IHC), Swine samples |
| Abcam, UK | Rb pAb anti-RUNX2 (ab23981) | WB (1:750) |
| Merk Millipore, UK | Rb pAb anti-Sox9 (AB5535) | WB (1:2,000) |
| Santa Cruz, Insight Biotechnology Ltd, UK | Ms mAb anti-osteocalcin (C-8, sc-74495) | WB (1:100) |
| Thermo Fisher Scientific, UK | Rb pAb anti-osteopontin (PA5-34579) | WB (1:1,000) |
| Abcam, UK | Rb mAb anti-Calumenin [EPR9075] (ab137019) | WB (1:500) |
| Abcam, UK | Rb pAb anti-GDF5 (ab23981) | WB (1:500) |
| Abcam, UK | Rb mAb anti-GLUT1 [EPR3915] (ab115730) | WB (1:10,000) |
| Merk Millipore | Rb pAb anti-ACVR1B | WB (1:1,000) |
| Santa Cruz, Insight Biotechnology Ltd, UK | Ms mAb anti-MeCP2 [G-6] (sc-SC-137070) | WB (1:750) |
| Novus Biologicals, Bio-Techne Ltd, UK | Rb pAb anti-METTL25 (NBP1-82105) | WB (1:750) |
| Cell Signalling Technology, UK | Rb mAb anti-p300 (D8Z4E, #86377) | WB (1:400) |
| Abcam, UK | Rb pAb anti-HB EGF (ab92620) | WB (1:500) |
| Santa Cruz, Insight Biotechnology Ltd, UK | Ms mAb anti-Smad7 (B-8) (sc-365846) | WB (1:1,000) |
| GE healthcare, Thermo Fisher Scientific, UK | Amersham ECL Ms IgG, HRP-linked whole Ab (NA931V)-secondary Ab | WB (1:5000) |
| GE healthcare, Thermo Fisher Scientific, UK | Amersham ECL Rb IgG, HRP-linked whole Ab (NA9340V)-secondary Ab | WB (1:5000) |

Ms, mouse; Rb, rabbit; mAb, monoclonal antibody; pAb, polyclonal antibody; HRP, horseraddish peroxidase; WB, western blot; IHC, immunohistochemistry

**Supplementary Table III. List of oligonucleotides for clonning.**

| **Target**  **[position in 3’UTR]** | **Sequence** |
| --- | --- |
| CALU |  |
| (NM_001130674) | Fw: 5'-AAACTAGCGGCCGCTGTTTGCGCTACTGAGACTGTTACTACG-3' |
| [position 155-162] | Rv: 5'-TCGACGTAGTAACAGTCTCAGTAGCGCAAACAGCGGCCGCTAGTTT-3' |
| SLC2A1/GLUT1 |  |
| (NM_006516.2) | Fw: 5'-AAACTAGCGGCCGCCAAAAGCAAGACTGTTGCTCAAATCTG-3' |
| [position 192-198] | Rv: 5'-TCGACAGATTTGAGCAACAGTCTTGCTTTTGGCGGCCGCTAGTTT-3' |
| GDF5 |  |
| (NM_001319138.1) | Fw: 5'-AAACTAGCGGCCGCAGTGTGAGGCTGTTAGACTGTTAGATG-3' |
| [position 508-515] | Rv: 5'-TCGACATCTAACAGTCTAACAGCCTCACACT GCGGCCGCTAGTTT-3' |
| ACVR1 |  |
| (NM_001105.4 ) | Fw: 5'-AAACTAGCGGCCGCGCATTCCTTACTTGCACTGTTACTCTG-3' |
| [position 452-458] | Rv: 5'-TCGACAGAGTAACAGTGCAAGTAAGGAATGCGCGGCCGCTAGTTT-3' |
| MECP2 |  |
| (ENST00000303391.6) | Fw: 5'-AAACTAGCGGCCGCACCAACAAGAATAAAGGCAGCTGG-3' |
| [position 43-49] | Rv: 5'-TCGACCAGCTGCCTTTATTCTTGTTGGTGCGGCCGCTAGTTT-3' |
| EP300 |  |
| (ENST00000263253.7) | Fw: 5'-AAACTAGCGGCCGCTTGGATCACTGTATAGACTGTTA-3' |
| [position 508-515] | Rv: 5'-TCGACTAACAGTCTATACAGTGATCCAAGCGGCCGCTAGTTT-3' |
| METTL25 |  |
| (NM_032230.2) | Fw: 5'-AAACTAGCGGCCGCCAAAAATTTTTAAATGACTGTTATGTG-3' |
| [position 43-49] | Rv: 5'-TCGACACATAACAGTCATTTAAAAATTTTTGGCGGCCGCTAGTTT-3' |

Fw, forward oligonucleotide; Rv, reverse oligonucleotide

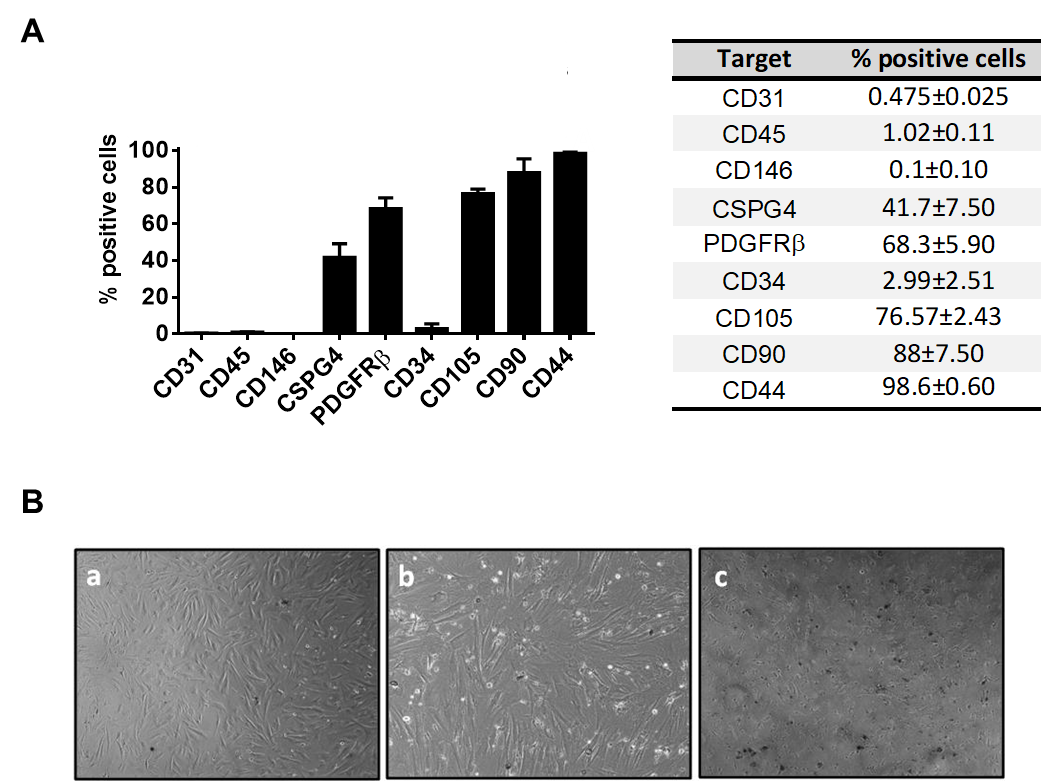

**A**

**B**

**Supplementary Figure 1: APCs and BM-MSCs characterization. A**, Validation of antigenic profile in culture-expanded APCs by flow cytometry**; B,** Effect of BM-MSC stimulation by high phosphate (HP) with or without high glucosa (HG) for 24h. **a**, Unstimulated BM-MSCs; **b**, BM-MSCs stimulated with HP; **c**, BM-MSCs stimulated with HG/HP causing dystrophic mineralization.

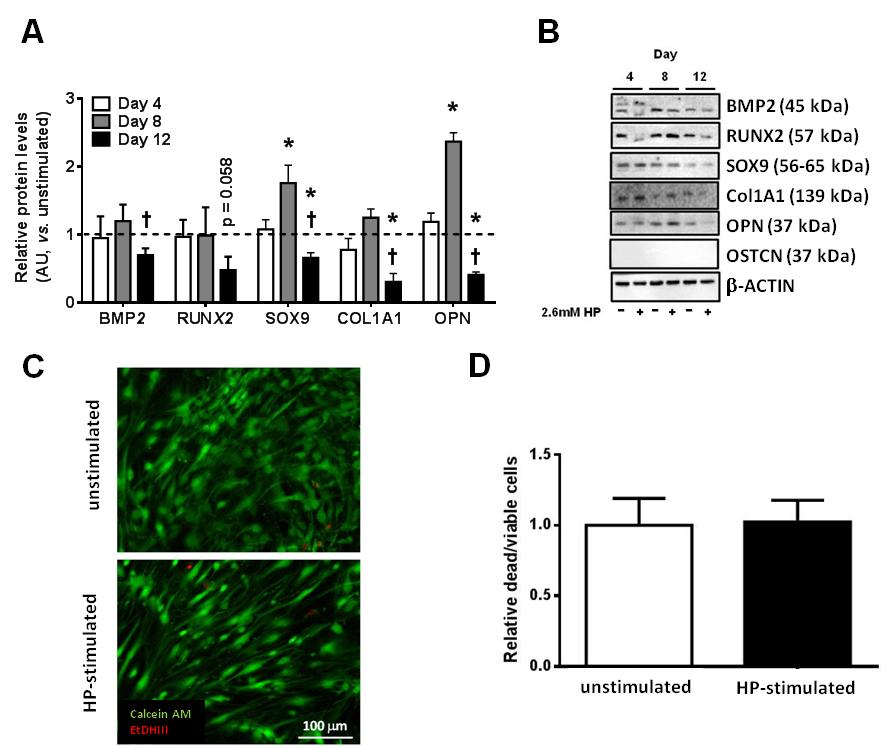

**Supplementary Figure 2: Effect of high phosphate on APC gene expression and viability**. **A,** Effect of high phosphate (HP) conditioning for 4, 8 and 12 days on osteoblast markers expression by APCs, as assessed by western blotting. Band densitometry was normalized to β-actin. Expression levels were standardized by basal values and are shown as arbitrary units (AU). **B,** Representative Western blotting of osteoblast markers. **C**, Representative fluorescence microphotographs showing viable (Calcein AM staining, green) and dead or poorly viable (ethidium homodimer-III/EtDHIII, red) APCs in unstimulated and HP-stimulated conditions. **D**, Bar graph showing viability data (relative dead/live cell counts). All experiments were performed in APCs from 4 (**A&B**) or 5 donors (**C&D**) in technical duplicates or triplicates. Plotted data are represented as mean ± SEM; *p<0.05 vs unstimulated; † p<0.05 vs day 8.

**Supplementary Figure 2: Effect of HP on APC gene expression and viability**. **A,** Effect of HP conditioning for 4, 8 and 12 days on osteoblast markers expression in APCs, as assessed by western blotting. Expression levels standardized by basal values. Band densitometry was normalized to β-actin. **B,** Representative western blotting of osteoblast markers. **C**, Representative fluorescence microphotograph showing viable (Calcein AM staining, green) and dead or poorly viable (ethidium homodimer-1, red) APCs in unstimulated and HP-stimulated conditions. **D**, Bar graph showing viability data (relative dead/live cell counts). All experiments were performed in APCs from 4 (**A&B**) or 5 donors (**C&D**) in technical duplicates or triplicates. Plotted data are represented as mean ± SEM; *p < 0.05.

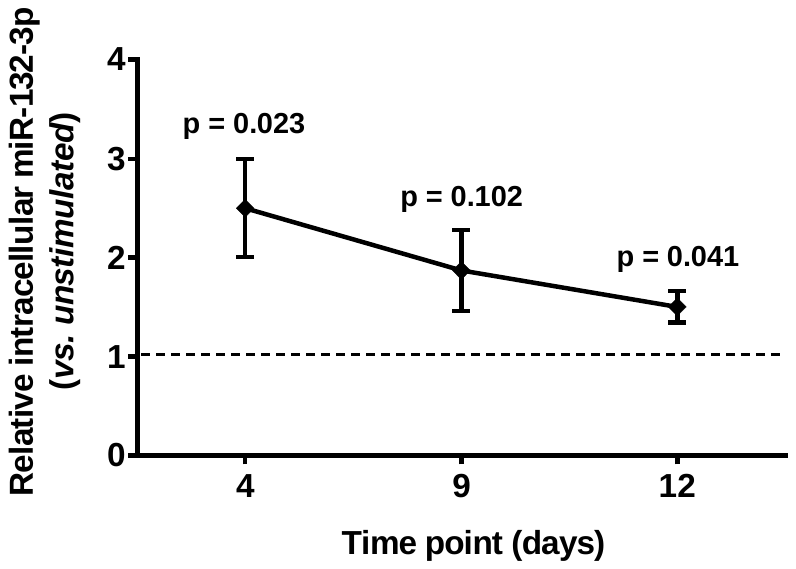

**Supplementary Figure 3: Expression of intracellular miR-132 at different time points following stimulation with high phosphate.** APCs from 5 donors were assessed. Plotted data are presented as mean ± SEM.

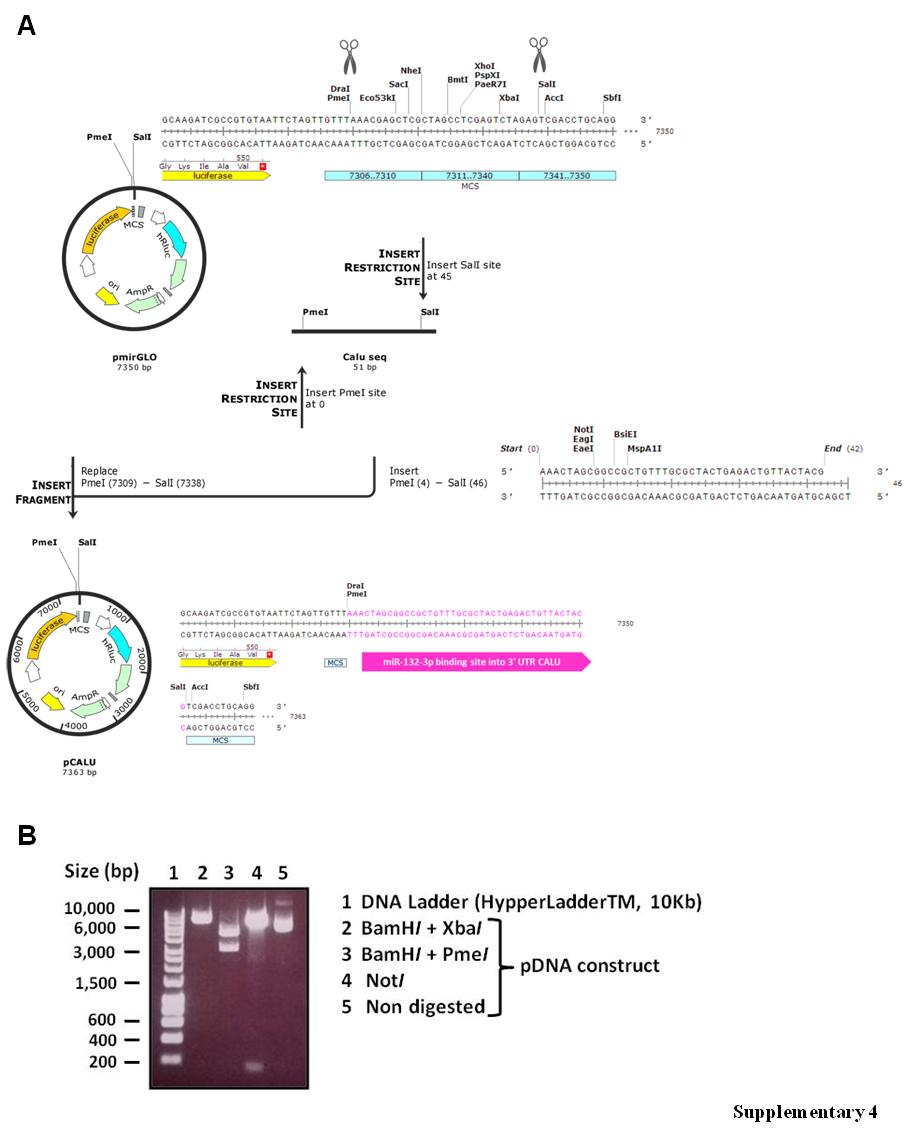

**Supplementary Figure 4. Preparation of miR132 target luciferase constructs. A**, SnapGene® drawing explaining cloning steps. **B**, Representative transilluminator picture of digestion products for pGLUT1 (pDNA) construct preparation to validate GLUT1 insert ligation into the expression vector pmiRGLO.

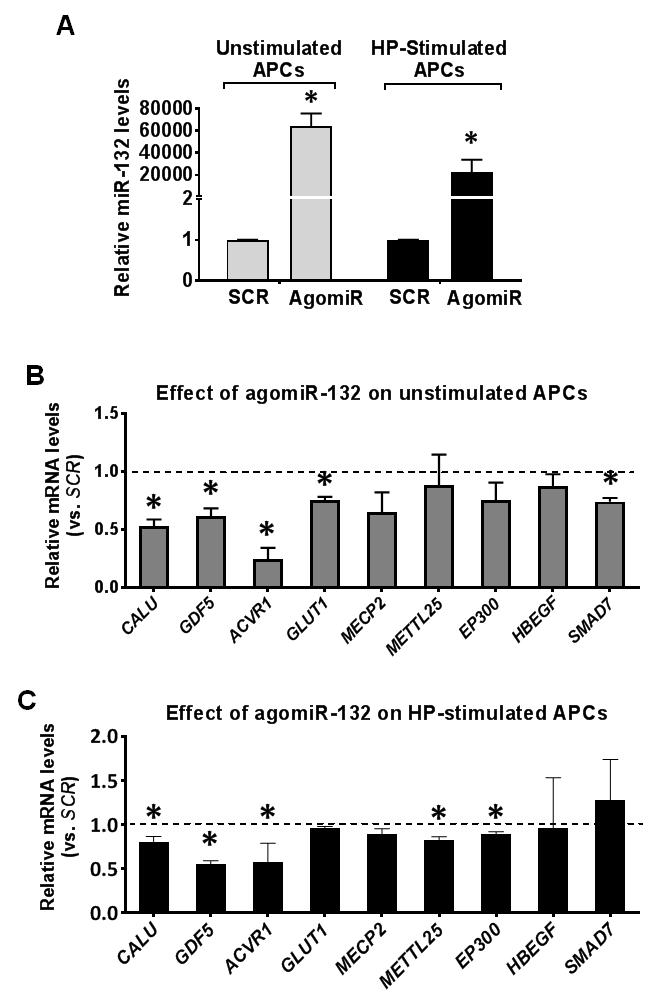

**Supplementary Figure 5. Effect of miR-132 forced expression on APC gene expression profile. A,** miR132 expression was transiently induced in APCs using agomiR either in unstimulated or HP-stimulated conditions. **B&C**, Effect of miR132 overexpression on predicted targets in unstimulated (**B**) and HP-stimulated APCs (**C**). All experiments were performed in APCs isolated from 4 different donors in technical triplicates. *p < 0.05 vs. Sramble (SCR).

**Supplementary Figure 5. Effect of miR-132 over-expression on APC profile. A**, miR132 expression was transiently overexpressed in APC (agomiR/miR mimic experiments) either in unstimulated or HP-stimulated conditions. **B&C**, Effect of miR132 overexpression on predicted targets in unstimulated (**B**) and HP-stimulated APCs (**C**). All experiments were performed in APCs isolated from 4 different donors intechnical triplicates. *p < 0.05 vs. Sramble (SCR).

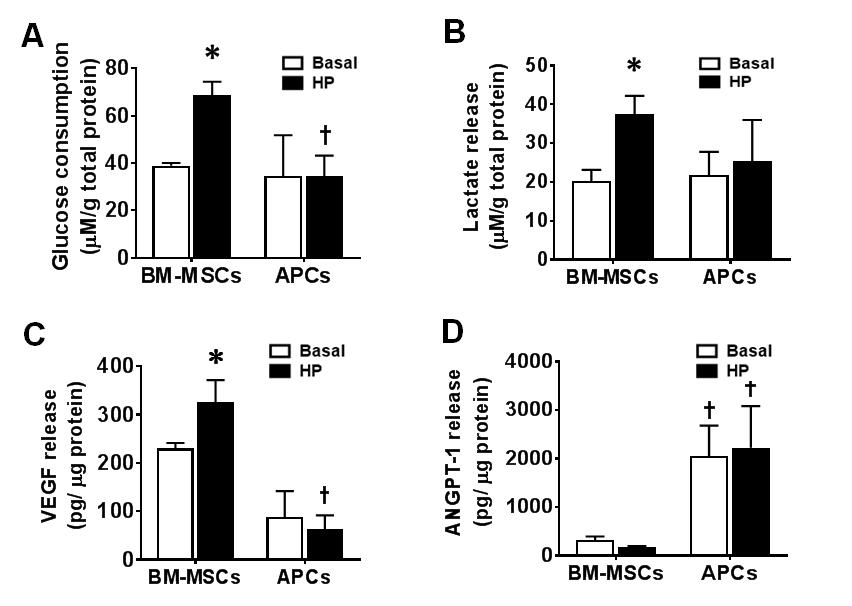

**Supplementary Figure 6. Effect of HP on metabolic features of APCs. A,** Effect of HP on glucose consumption in APCs and BM-MSCs. **B,** Effect of HP on lactate release. **C,** Effect of HP on VEGFA release assessed by measuring levels in CCM. **D,** Effect of HP on ANGPT1 release. APC isolated from 3 donors, assayed in technical triplicates per condition. Values are mean ± SEM. *p < 0.05 vs. respective control, ^†^P<0.05 vs. BM-MSCs.

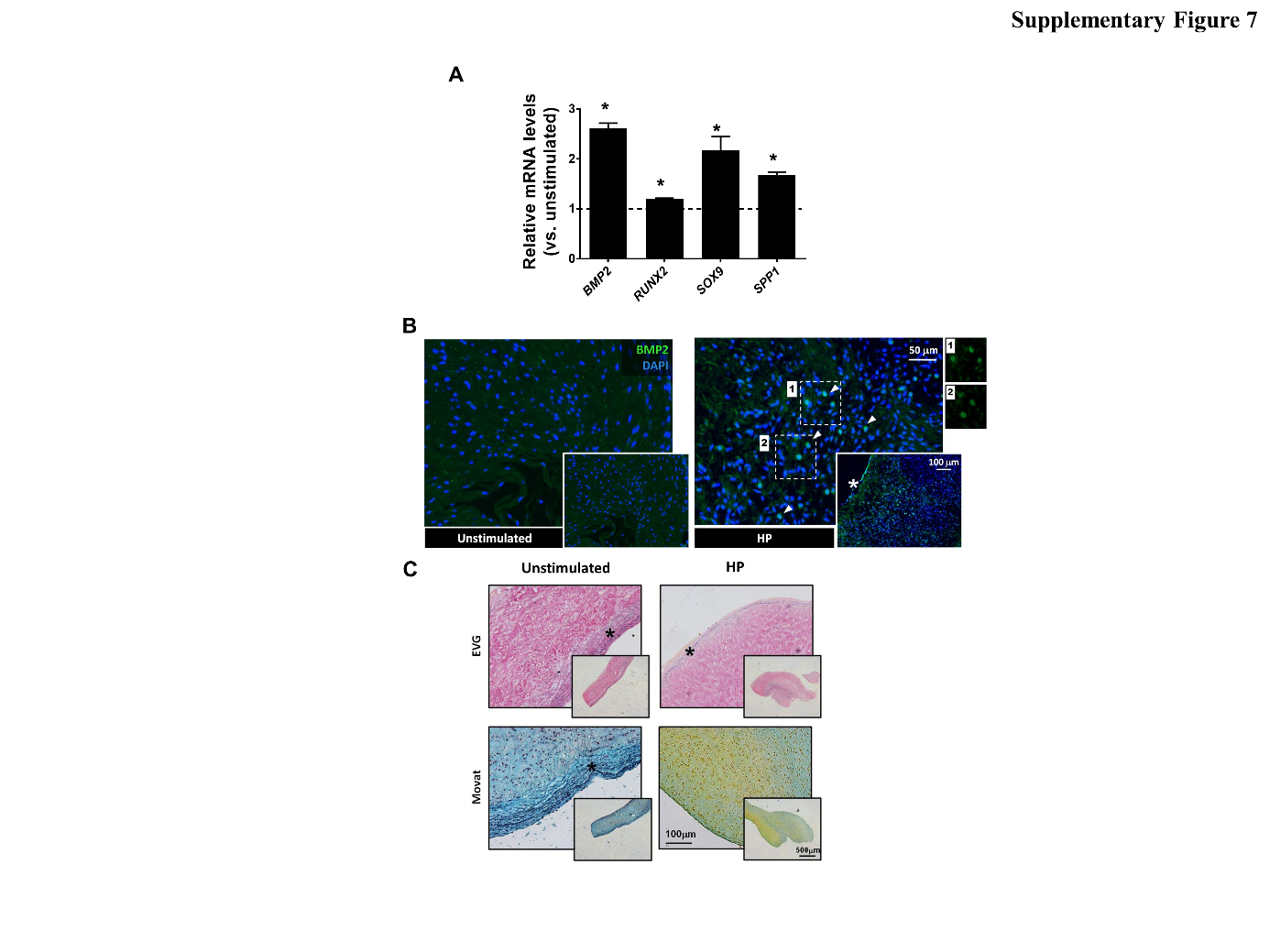

**Supplementary Figure 7. Explanted swine aortic valve assay.** **A,** Changes in gene expression after exposure of EAVs to high phosphate (3.2 mM). Data are represented as mean ± SEM; *p<0.05 vs. unstimulated. **B,** Immunofluorescence images of BMP2 expression. (*) indicates areas of calcification. **C,** Staining for elastin (EVG) and collagen (Movat). (*) in EVG and Movat indicate elastin deposition
